# TRESK background K^+^ channel deletion selectively uncovers enhanced mechanical and cold sensitivity

**DOI:** 10.1101/636829

**Authors:** Aida Castellanos, Anna Pujol-Coma, Alba Andres-Bilbe, Ahmed Negm, Gerard Callejo, David Soto, Jacques Noël, Nuria Comes, Xavier Gasull

## Abstract

Changes in TRESK K^+^ channel expression/function enhance sensory neurons excitability, but its role in somatosensory perception and nociception is poorly understood. We show that TRESK regulates the sensitivity to mechanical and cold stimuli but not the perception of heat. TRESK knockout mice nociceptive neurons present an enhanced excitability; skin nociceptive C-fibers show an increased activation by lower intensity cold or mechanical stimulation and mice lacking TRESK present mechanical and cold hypersensitivity. TRESK is also involved in osmotic pain and in early phases of formalin-induced inflammatory pain, but not in the development of mechanical and heat hyperalgesia during chronic pain. In contrast, mice lacking TRESK present cold allodynia that is not further enhanced by oxaliplatin. In summary, genetic removal of TRESK uncovers enhanced mechanical and cold sensitivity, indicating that it regulates the excitability of specific neuronal subpopulations involved in mechanosensitivity and cold-sensing, acting as a brake to prevent activation by low-intensity stimuli.

## Introduction

Sensory perception is triggered by excitation of sensory neurons terminals in the periphery and generation of action potentials that carries sensory information towards the central nervous system. The combined activation of different ion channels and membrane receptors determine the likelihood of excitation and generation of action potentials in specific subtypes of sensory neurons such as thermoreceptors, mechanoreceptors or nociceptors. Two-pore domain potassium channels (K_2P_) are expressed in different subpopulations of sensory neurons, including nociceptors where they carry most of the “leak” or background current [1]. Their electrophysiological properties allow them to carry K^+^ currents over a wide range of membrane potentials and hence they are key determinants of neuronal excitability, decreasing the probability of depolarizing stimuli to achieve action potential threshold, as well as shaping the neuron firing response [1]. TRESK, TREK-1, TREK-2 and TRAAK channels make the major contribution to leak currents in trigeminal (TG) and dorsal root ganglion (DRG) neurons [2-6], where they have been implicated in perception of pain induced by mechanical, thermal and chemical stimuli, as well as in neuropathic and inflammatory pain [4,7-11]. Depending on the specific expression and properties of each one of the K_2P_ channels in different subpopulations of sensory neurons they fine-tune the sensitivity of neuronal subpopulations to noxious stimuli. In this regard, deletion of one of the K_2P_ channels enhances the specific sensitivity to certain external stimuli but not others, rather than acting as a general effect on neuronal excitability due to the removal of hyperpolarization by K^+^ currents. TRESK shows a high expression in DRG and TG sensory neurons in human, rat and mouse and is particularly enriched in sensory ganglia compared to other neural and non-neural tissues [3,5,10,12-16]. Single-cell RNA sequencing data has provided further insight on the potential role of TRESK in sensory neurons, since the channel is predominantly expressed in non-peptidergic nociceptors (NP1-NP3) as well as in a subpopulation of low-threshold mechanoreceptors, while its expression is lower in peptidergic nociceptors [15,16]. Interestingly, TRESK expression in DRG neurons is downregulated in different pain conditions, comprising sciatic nerve axotomy [4], spared nerve injury [17] and chronic inflammation [10], which contributes to the enhancement of neuronal excitability. In contrast, different experimental manipulations to increase TRESK expression resulted in a decrease in neuronal excitability and amelioration of painful behaviors [17-19]. In the same line of evidence, sensory neurons from a functional TRESK[G339R] knockout mice showed a significant reduction of outward K^+^ current, increased excitability and reduced rheobase [3]. Also, a frameshift mutation leading to the truncation of the channel has been associated with familial migraine with aura, thus involving the channel in the enhanced activation of the trigeminovascular system and the release of inflammatory neuropeptides (CGRP/substance P) in the meninges and cerebral vessels triggering pain associated to migraine [20]. Recent studies show that, to trigger migraine pain, not only TRESK malfunction is needed but also the combined down-regulation of TREK-1/2 induced by the heteromerization with truncated TRESK proteins that prevent membrane channel expression [21]. Despite the fact that TRESK shares some functional properties with other K_2P_ channels, its high expression in sensory ganglia and its selective expression in specific subsets of sensory neurons points to a relevant role of this channel in sensory perception and pain that is not yet well understood. Here, we report that removal of TRESK uncovers enhanced cold and mechanical sensitivity without affecting thermal sensitivity to warmth or hot temperatures.

## Results

### TRESK deletion reduces total background current and enhances neuronal excitability

TRESK channels have been detected in small- and medium-sized sensory neurons, together with other members of the K_2P_ family of background K^+^ channels [5,15,16]. To confirm the elimination of TRESK expression in homozygous knockout animals and to assess possible effects on the expression of other K_2P_ channels induced by knocking out TRESK, we first studied the mRNA expression in DRGs by real-time quantitative PCR. TRESK mRNA was undetected in KO mice while a significant expression was found in wild-type (WT) animals (Fig 1A). Since removal of TRESK could potentially induce compensatory effects on other K_2P_ channels involved in pain perception, the expression of TREK-1, TREK-2 and TRAAK was determined (the more expressed K_2P_s in sensory neurons, together with TWIK1, which is unclear if it forms functional membrane channel). Analogously to what has been reported in a previous study [22], mRNA for these channels were present in WT and KO mice at similar levels, thus excluding a compensation effect on the expression of these K_2P_ channels in KO mice. Interestingly, the expression of TRPA1 and TRPV1, two channels highly expressed in nociceptors, was not significantly changed.

**Figure 1:**
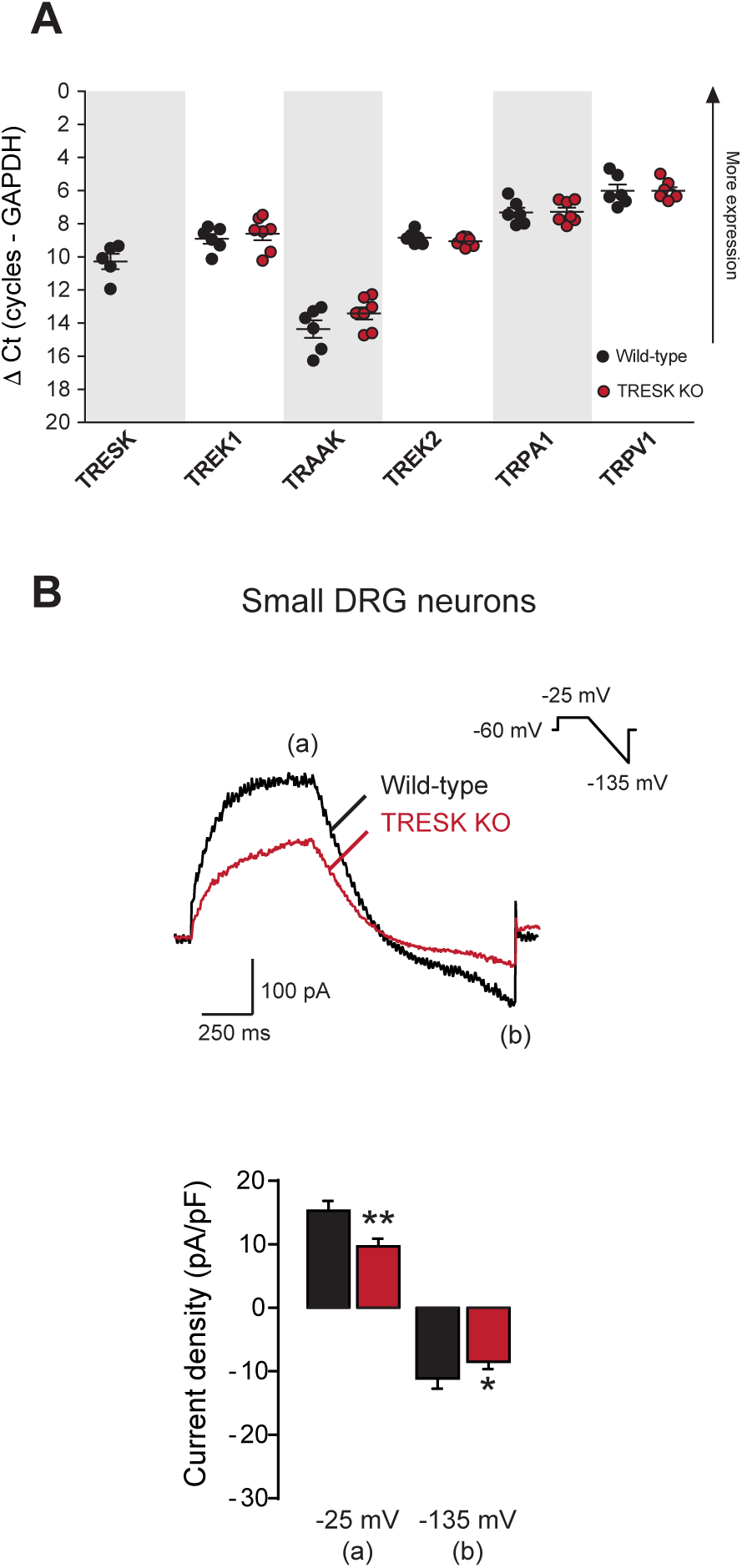
Nociceptive sensory neurons lacking TRESK have a decreased standing outward current. **A.** Expression profile of K_2P_ channels, TRPA1 and TRPV1 in mouse sensory neurons from wild-type and TRESK knockout mice. mRNA expression obtained by quantitative PCR shows no significant differences between WT and KO animals after genetic deletion of TRESK. Y-axis shows the ΔCt (number of cycles target - number of cycles GADPH) for the different mRNAs. Notice that the Y-axis has been inverted to visually show that lower ΔCt numbers are indicative of a higher expression. Each dot represents a single animal (wild-type n=6; TRESK knockout n=7). **B.** Top: Representative recordings of whole-cell currents from small-sized DRG sensory neurons using a protocol to minimize activation of voltage-gated transient K^+^ outward currents (holding voltage - 60mV). Bottom: quantification of currents at the end of the pulse at −25 mV (a) and at the end of the ramp (−135 mV; b) showed significant differences among groups (*p<0.05; **p<0.01 unpaired t-test. wild-type n=31; TRESK KO n=50).

To assess whether removal of TRESK modifies K^+^ background currents, we next recorded total K^+^ current from small diameter DRG neurons, that are usually assumed to be nociceptive neurons (soma size <30 μm; capacitance <30pF). For this, as previously described [3,4,23,24], we measured the current density at −25 mV followed by a voltage ramp to −135 mV (Fig 1B). Wild-type sensory neurons presented a significantly larger current density compared to KO neurons both at −25 mV (for WT, 15.3±1.5; n=31; Cm=21.7±1.8 pF; for KO, 9.7±1.2 pA/pF; n=50; Cm=22.1±1.0 pF; unpaired t-test p=0.004), and −135 mV (for WT, −11.1±1.6 pA/pF; for KO, −8.5±1.2 pA/pF; unpaired t-test p=0.038), indicating that the absence of TRESK has a significant functional consequence in the membrane current. In a second batch of recordings from small-sized DRG neurons, we investigated the functional effects of TRESK removal on neuronal excitability by measuring their resting membrane potential and action potential properties (Fig 2A and B). Wild-type and TRESK KO sensory neurons did not present significant changes in resting membrane potential (RMP:−58.3±3.9 vs. - 56.2±1.5 mV, respectively. Unpaired t-test p=0.540; Fig 2B), suggesting that TRESK might have a minor role in setting the RMP and other K^+^ channels are possibly more important in setting the RMP in the cell body. This is in agreement with previous data where RMP was not significantly modified after TRESK down-regulation or deletion [3,4]. Interestingly, injected current threshold to action potential firing in current-clamp configuration was significantly decreased in TRESK KO neurons (Fig 2B, p=0.0013 unpaired t-test) compared to neurons from WT littermates. This effect is likely a consequence of the increased membrane resistance found in TRESK KO neurons (1262.8±133.8MΩ) compared to controls (Wild-type: 792.7±186.9MΩ; p=0.045 unpaired t-test; Fig 2B). As expected, action potential amplitude was not significantly altered (p=0.406, unpaired t-test) since this parameter is more dependent on voltage-dependent sodium channels (Na_v_) activity. However, action potentials were significantly wider in neurons from TRESK KO, likely reflecting the consequence of the decrease in total K^+^ current (50% AP width TRESK KO 4.03±0.5 s; n=27 neurons; Wild-type: 2.38±0.5 s; n=13; p=0.041; Fig 2B). To investigate if these changes affect the excitability of sensory neurons, we injected a current ramp (0 to 500 pA, 1s) in current-clamp configuration and counted the number of action potentials fired for each genotype. As shown in Fig 2C, TRESK KO neurons fired on average more spikes (9.0±1.3 spikes) than wild-type neurons (5.5±1.1 spikes; p=0.036), indicating that removal of TRESK increased the excitability of sensory neurons.

**Figure 2:**
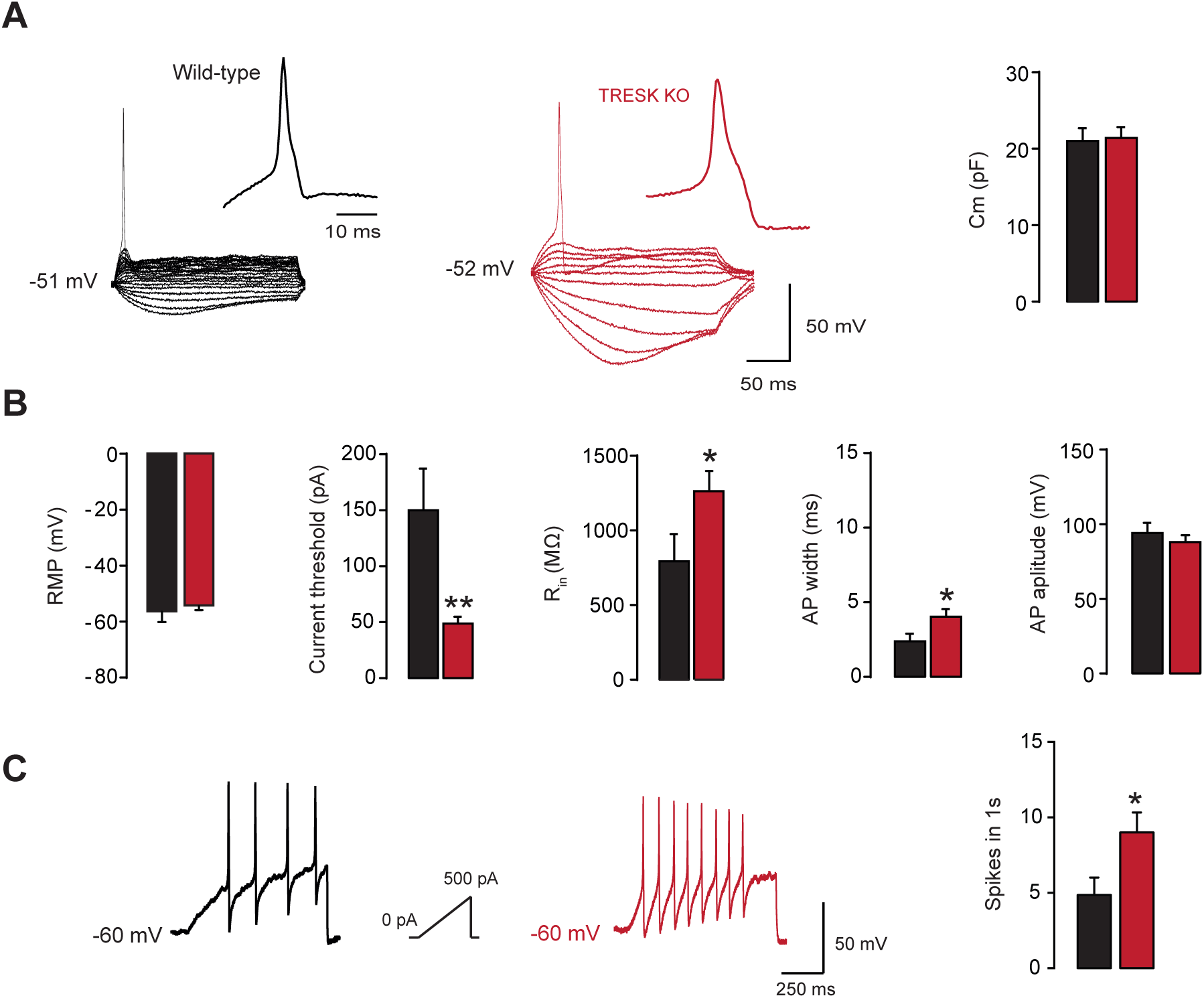
Nociceptive sensory neurons lacking TRESK present a higher excitability. **A.** Representative whole-cell current-clamp recordings from wild-type and TRESK knockout nociceptive sensory neurons elicited by hyperpolarizing or depolarizing 400 ms current pulses in 10 pA increments. *Right*: mean membrane capacitance (C_m_) from neurons studied is shown. **B.** Quantification of the electrophysiological parameters analyzed in wild-type (black bars, n=13) and TRESK KO (red bars, n=27) sensory neurons. RMP: resting membrane potential. Current threshold was measured using 400 ms depolarizing current pulses in 10 pA increments; R_in_: whole-cell input resistance was calculated on the basis of the steady-state I-V relationship during a series of 400 ms hyperpolarizing currents delivered in steps of 10 pA from −50 to −10 pA. AP amplitude was measured from the RMP to the AP peak. AP duration/width was measured at 50% of the AP amplitude. Statistical differences between groups are shown (*p<0.05, **p<0.01 unpaired t-test). **C.** Examples and quantification of neuronal excitability as the number of action potentials fired in response to a depolarizing current ramp (0 to 500 pA, 1s) form a holding voltage of −60 mV (black, wild-type n=6; red, TRESK KO n=7). Statistical differences between groups are shown (*p<0.05; **p<0.01 unpaired t-test).

To further characterize the responsiveness of sensory neurons in TRESK KO mice, we measured the intracellular Ca^2+^ signals ([Ca^2+^]) of cultured DRG neurons from WT and KO mice in response to capsaicin (1 μM; TRPV1 agonist) and AITC (100 μM; TRPA1 agonist), two markers of nociceptive neurons, and menthol (100 μM; TRPM8 agonist), a marker of cold sensory neurons. Among wild-type DRG neurons (total number of neurons analyzed = 1124), 49.2% responded to capsaicin, 40.4% to AITC and 7.0% to menthol. Responses to both capsaicin and AITC were seen in 17.9% of the neurons (Fig 3A and B). A percentage of neurons did not respond to any of the agonists tested (246 neurons, 21.9%). Responses to capsaicin (45.7%) and menthol (9.1%) were not different in neurons from TRESK KO mice (n=1228) but, the percentage of neurons activated by AITC was significantly lower (24.7%, p=0.009). Neurons non-responding to any agonists were 446 (36.3%). The diameters of sensory neurons responding to the different agonists between WT and KO animals were not different which means that they represent identical populations of DRG neurons (mean soma diameter WT: 20.2±0.19 μm; KO: 20.3±0.23 μm). In parallel experiments, the response of cultured trigeminal neurons was assessed. These showed no significant differences for the response to the different agonists (data not shown) between genotypes. Therefore, the diminished response to AITC is restricted to DRG neurons.

**Figure 3:**
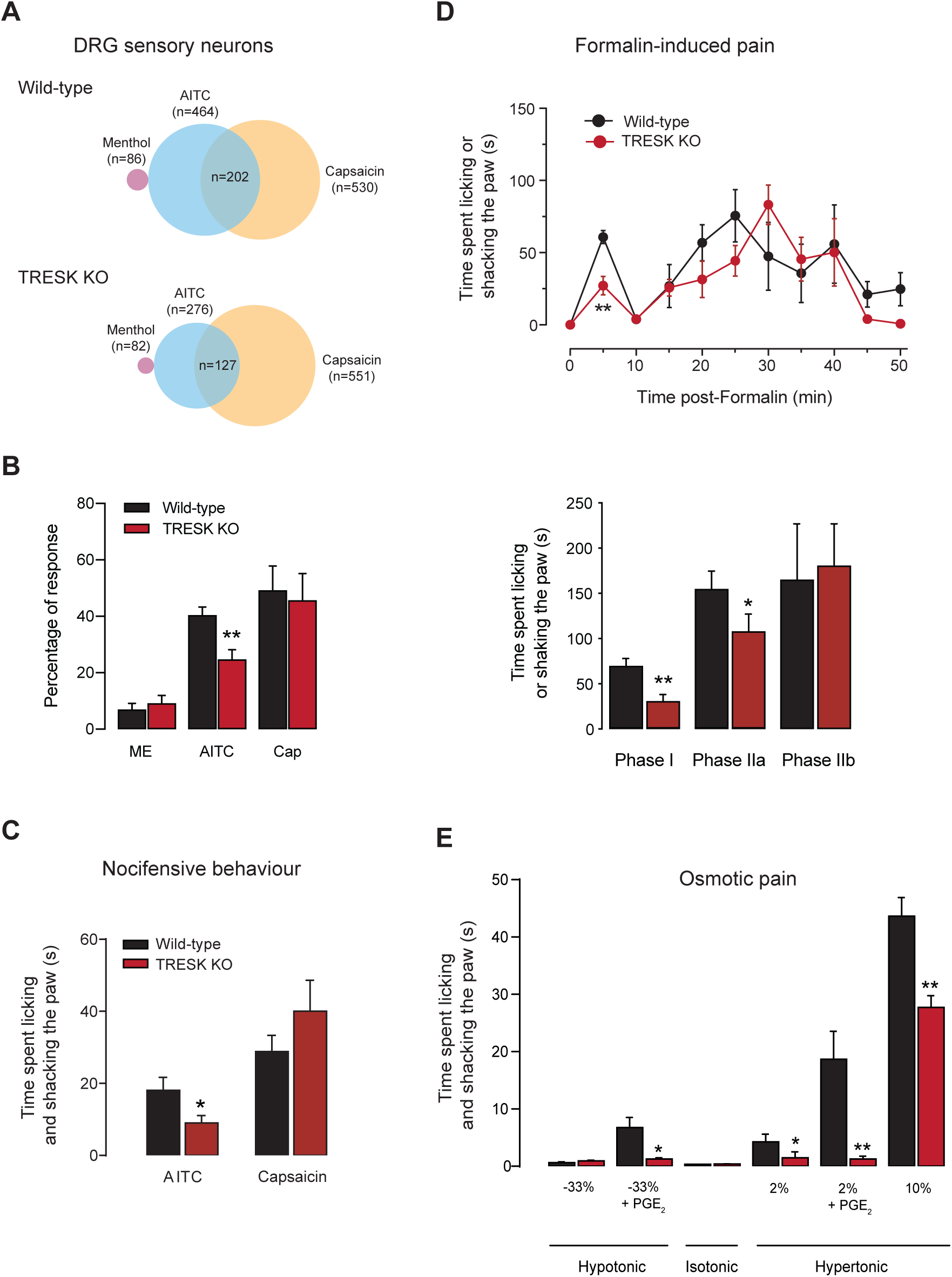
TRESK KO mice present diminished responses to osmotic stimuli and TRPA1 activation. **A.** Venn diagrams showing the relative size and overlap between the population of neurons activated by menthol (100 μM), allyl isothiocyanate (AITC, 100 μM) and capsaicin (1 μM) using Fura-2 ratiometric intracellular calcium imaging. Number of total cells analyzed was 1124 (wild-type) and 1228 (knock-out). The number of responding cells in each subgroup is shown in parenthesis. Non-responding neurons to any of the agonists assayed was 246 (wild-type) and 446 (knock-out). **B.** Quantification of the percentage of neurons responding to each agonist in intracellular calcium recordings. Statistical differences between groups are shown (**p<0.01 unpaired t-test). **C.** Nocifensive behavior: quantification of the time the animals spent licking and shaking the paw after intradermal injection of AITC (10%) or capsaicin (1μg/10μl) in the hind paw for wild-type and TRESK knockout mice. The mean time spent showing nocifensive behaviors over a period of 5 min is shown (WT n=13; KO n=13 animals). **D.** Formalin-induced pain: *Top*. Time course of licking/shaking behavior directed to the formalin-injected hind paw. *Bottom*: quantification of cumulative time spent in phase I (0-10 min), IIa (11-30 min) and IIb (31 to 50 min). n= 5-6 animals per group. **E.** Osmotic pain: quantification of the time spent licking and shaking the paw after intradermal injection of hypotonic (−33%) or hypertonic (2% and 10%) stimuli in the hind paw of wild-type and TRESK knockout mice. The mean time spent showing nocifensive behaviors over a period of 5 min is shown. 6 to 11 animals were evaluated in each group. When indicated, a 5 μl-injection of prostaglandin E_2_ (PGE_2_) was injected in the hind paw of each mouse 30 min before the test to sensitize nociceptors. Statistical differences are shown as *p<0.05, **p<0.01 (Student’s unpaired t-test) between WT and KO mice.

We next assessed if the reduced response to AITC of DRG neurons from TRESK KO mice had a behavioral correlate by injecting AITC (100 μM) into the mouse hind paw. In agreement with the previous observation, nocifensive behavior measured as the time spent licking or shaking the paw significantly diminished in KO compared to WT mice (TRESK KO: 9.0±2.4 s; WT: 18.1±3.6 s; p=0.039; n=13 for each group; Fig 3C). In contrast, painful behavioral responses to capsaicin injection did not differ between KO and WT mice (40.0±8.6s vs. 28.8±4.5s, respectively; p=0.236; Fig 3C). Injection of vehicles for each compound did not produce significant behavioral effects (data not shown). To further investigate to what extend chemical nociception was altered by the absence of TRESK, we used the formalin test [25,26], which is characterized by an initial phase (phase I; 0-5 min) due to direct nociceptor activation [27] and a second phase (phase II; 15-50 min) that is attributed to a combined nociceptive input together with central spinal sensitization [26]. The phase I nocifensive response has been directly linked to the activation of TRPA1 [27] although high concentrations of formalin are still able to induce some pain in TRPA1 KO mice [28]. Interestingly, animals lacking TRESK showed a diminished response to formalin injection in phase I (30.8±6.1s; p=0.002; Fig 3D) compared to control WT animals (69.9±7.2s). The decreased response in phase I of TRESK KO animals is in agreement with the decreased nocifensive response observed after AITC injection (Fig 3C) which corroborates the decreased fraction of AITC sensitive DRG neurons from TRESK KO in culture. All these observations confirm that TRPA1 activation is decreased in TRESK KO animals. Phase II of the formalin test can be further split in phase IIa (15-25 min) and IIb (25-50 min), where IIa has a higher nociceptive input than IIb. TRESK KO animals showed a decreased licking time during phase IIa (107.7±16.3s) compared to wild-type animals (154.8±18.9s; p=0.048); which probably reflects a decreased nociceptors activation as found in phase I. In contrast, phase IIb of the formalin test did not show significant differences and licking behavior was similar between groups (p=0.421, Fig 3D). Injection of hypertonic saline stimulates primary afferent nociceptors and produce pain in humans [29,30]. This response can be further enhanced by sensitization of nociceptors with PGE_2_. Injection of hypotonic stimuli did not induce significant nocifensive behavior in resting conditions (p=0.146, Fig 3E) but, as previously described [7,30], nocifensive responses were enhanced after sensitization with PGE_2_ in wild-type mice but not in TRESK KO mice (p=0.019, unpaired t-test WT vs. KO). Nocifensive responses to mild hypertonic saline were diminished in TRESK KO animals both, in resting conditions (2% or 10% NaCl, p=0.049 and p=0.002) or after sensitization with PGE_2_ (p=0.002; Fig 3E). Again, these effects are similar to the one reported after knocking out TREK-1 and TREK-2, but not in the single TRAAK knockout mice [7-9], thus implying that TREK-1, TREK-2 and TRESK are involved in the sensitivity to hypertonic stimuli and their absence prevents sensitization by PGE_2_.

### Mice lacking TRESK present mechanical allodynia and normal heat perception

Since TRESK knockout enhances nociceptive sensory neuron excitability, we next analyzed the sensitivity of nerve fibers from TRESK KO mice to different types of stimuli and whether the activation of sensory fibers was correlated to behavioral responses to these stimuli. Saphenous nerve C-fibers from WT (n=14) and TRESK KO mice (n=22) were recorded with the nerve-skin preparation. Mechanical thresholds were determined with calibrated von Frey filaments applied on the receptive fields of C-fibers. TRESK KO mice presented an enhanced sensitivity to mechanical stimuli. Mean threshold value for the whole population was not significantly different in KO animals (27.6±5.2 mN) compared to WT mice (33.2±6.2 mN; p=0.268 unpaired t-test), but threshold values from KO mice presented a wider distribution and a significant shift towards lower values (Fig 4A). Further analysis showed that KO mice presented a significant percentage of fibers with lower thresholds (<12 mN; 40.9%) compared to fibers from WT animals (0%; p=0.006; Fisher’s exact test). This suggests that TRESK function prevents C-fiber activation by low intensity mechanical stimuli and therefore deletion of the channel reveals mechanical sensitivity with low threshold values in a fraction of C-fibers (Fig 4A). We then compared the mechanical sensitivity of TRESK KO to WT mice with the von Frey up and down method. Hind paw mechanical sensitivity was significantly higher in TRESK KO mice compared to WT (p<0.0001 for males and females; Fig 4D), thus showing that lower C-fibers thresholds values produce mechanical allodynia in behaving mice. Measures of mechanical sensitivity with the dynamic plantar aesthesiometer also confirmed significant differences to pressure application between WT (5.06±0.12 g; n=27) and KO animals (4.66±0.17 g; n=22; t-test p=0.027; Fig 4D).

**Figure 4:**
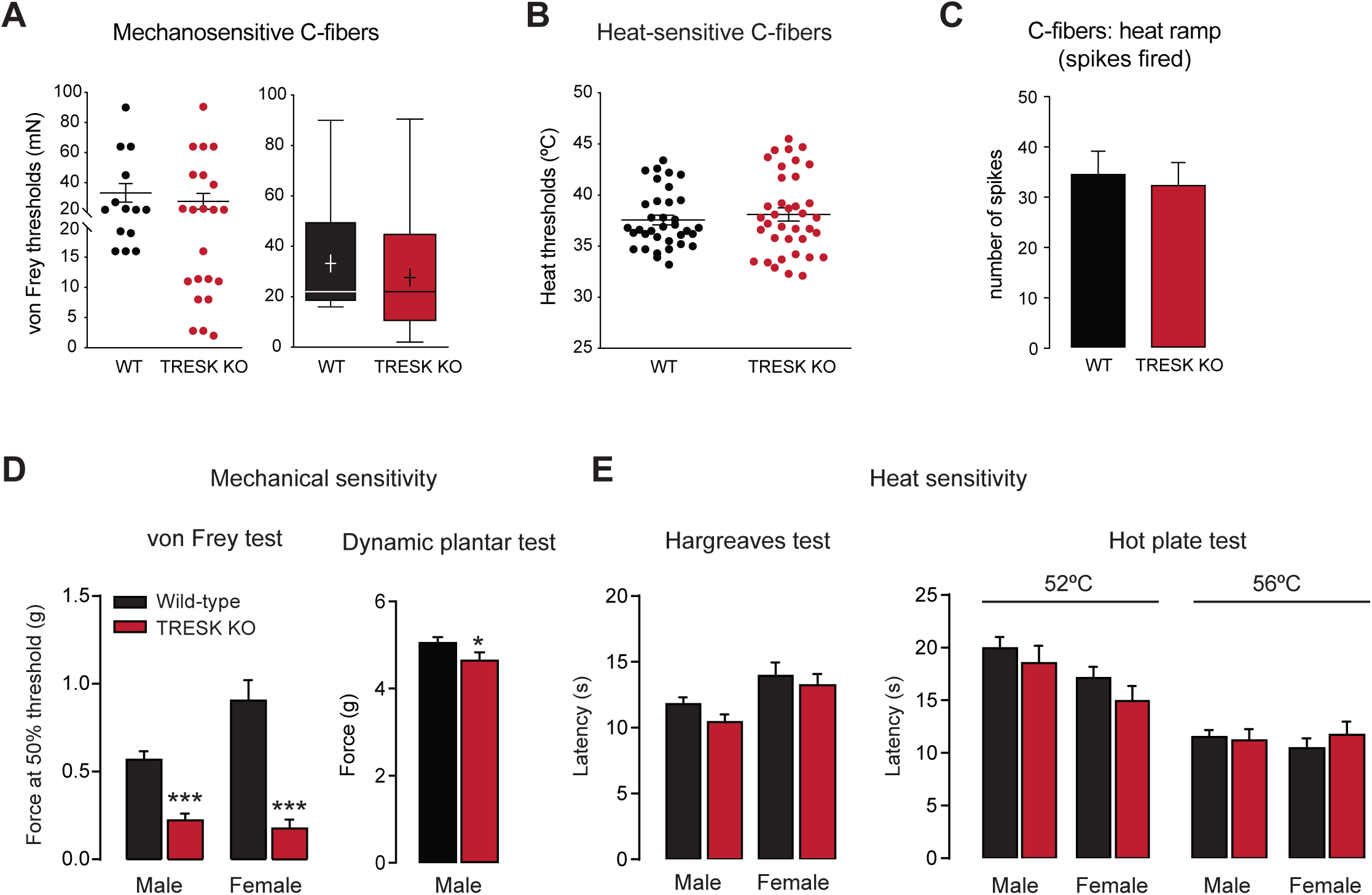
TRESK-lacking mice present mechanical allodynia and normal heat sensitivity. **A.** Mechanical threshold obtained with von Frey hairs from saphenous nerve C-fibers. *Left.* Distribution of von Frey thresholds. *Right*. Box and whiskers plot of the distribution of von Frey thresholds. Whiskers show the minimum and maximum values obtained. Mean value is shown as an + and median as a straight line. Wild-type n=14, knock-out n=22. **B.** Distribution of heat thresholds from C-fibers recorded from wild-type (n=35) and TRESK KO mice (n=37), measured in skin-nerve experiments. **C.** Mean number of spikes fired by heat sensitive C-fibers during a heat-ramp from 30 to 50°C. No significant differences were found. **D.** Mechanical sensitivity. *Left*: von Frey response thresholds obtained with the up and down method in male and female wild-type and TRESK knockout animals (male WT n=58, KO n=33; female WT n=18, KO n=19). *Right*: latency to hind paw withdrawal in the dynamic plantar test (male WT n=27; KO n=22). Statistical differences are shown as *p<0.05, ***p<0.001 (Student’s unpaired t-test) between WT and KO mice. **E.** Heat sensitivity. *Left*: radiant heat (Hargreaves test) in male (WT n=39; KO n=38); and female animals (WT n=18; KO n=15). *Right*: Noxious heat sensitivity to 52 and 56°C (Hot plate test) in male (WT n=8/9; KO n=13/11); and female animals (WT n=9/9; KO n=12/8).

Other channels of the K_2P_ family have been involved in thermosensation and pain perception in response to hot or cold stimuli [7-9]. TRESK activity is not modulated by changes in temperature in the physiological range [5]. Responses of C-fibers activated by heat-ramp applied on their receptive fields in the skin from TRESK KO and WT mice did not show significant differences in heat thresholds for the activation of C-fibers (37.6±0.46°C, n=35 C-fibers for WT; 38.1±0.64°C, n=37 C-fibers for KO; p=0.502 t-test; Fig 4B), nor the distribution of these thresholds in a range of temperatures, nor the number of spikes fired by heat sensitive C-fibers during heat-ramps (32.6±4.4 spikes WT; 34.8±4.4 spikes TRESK KO; p=0.720 t-test; Fig 4C), indicating that TRESK does not seem to have a major role in the detection of warm or hot temperatures by C-fibers. We then tested heat sensitivity of mice with Hargreaves and hot plate tests. Heat sensitivity was unaltered in TRESK KO mice compared to WT in the radiant heat Hargreaves test (p=0.08 males; p=0.603 females, Fig 4E), which measures the threshold sensitivity to heat confirming the observations made in C-fibers recordings. To further evaluate a possible implication of TRESK in heat pain, we evaluated the sensitivity to more extreme temperatures in the hot plate test at 52 and 56°C, but no significant differences were detected (Fig 4E). This indicates that the extreme heat sensitivity is conserved in TRESK KO mice. The dynamic hot plate with a ramp of temperature has been proposed as a valuable method to differentiate between thermal allodynia and hyperalgesia [31]. The number of jumps of the mice on a hot plate when the plate temperature is increased from 39 to 50 °C did not show any difference between TRESK KO and WT, neither for the total number of jumps nor the temperature at which animals elicited their first jump (Suppl. Fig 1). Only a small difference was found for TRESK KO and WT female mice at the high temperature of 49°C, where KO animals seem less sensitive, but not at 48 or 50°C. Both recordings of C-fiber activity and behavioral tests discard a significant contribution of TRESK in heat sensitivity.

### Elevated perception of cold temperatures in the absence of TRESK channel

Thermosensitivity to cold temperatures is governed by different mechanisms and by distinct neuronal subpopulations than warm/hot perception [32]. We evaluated the role of TRESK in the cold sensitivity of C-fibers by recording fibers activity with the saphenous nerve preparation upon cooling their receptive fields in the skin from 30 to 10°C over 90 s (Fig 5A-D). C-fiber recordings from TRESK KO mice showed a significant change in the fractions of mechano-cold and mechano-heat and cold sensitivity of polymodal C-fibers compared to WT mice. TRESK KO increased the fraction of mechano-cold C-fibres compared to WT mice (37% and 20% C-MC, respectively; Fig 5A). This was accompanied by a reduction of the percentage of mechano-heat-cold C-fibers (C-MHC) in KO mice (29%) compared to WT (38%; p<0.01; Chi-square test; Fig 5A). Nevertheless, the total number of cold sensitive fibers did not differ significantly between genotypes (KO: 66% of C-fibers; WT: 58%). The distribution of cold C-fibers thresholds showed a clear shift towards warmer temperatures, between 26 and 28°C, in the TRESK KO, while fiber thresholds from WT animals were more uniformly distributed at different temperature segments (Fig 5B and 5C). Despite mean cold thresholds were not different (p=0.388, Fig 5C), we could observe a tendency to have more fibers activated at higher temperatures above 25°C in knockout animals (58%) compared to WT (42%), although this difference did not reach significant statistical power (Fisher’s exact test; p=0.099). The total response of TRESK KO fibers to the cooling of 90 s (33.1±5.1 spikes) was similar to WT (34.1±5.8 spikes; p=0.90 unpaired t-test), however, the distribution of the activity upon cooling was higher at temperatures between 25 and 19°C for TRESK KO fibers (Fig 5D). The baseline activity between 30-31°C was also higher in TRESK KO than WT fibers (0.3 spikes/s for TRESK KO and 0.1 spikes/s for WT; n=36 and n=31 respectively; p<0.001, t-test). These experiments indicate that the temperature for activation of cold-sensitive C-fibers was increased in TRESK KO, but also that the overall fiber activity in response to cooling did not depend drastically on the presence of TRESK.

**Figure 5:**
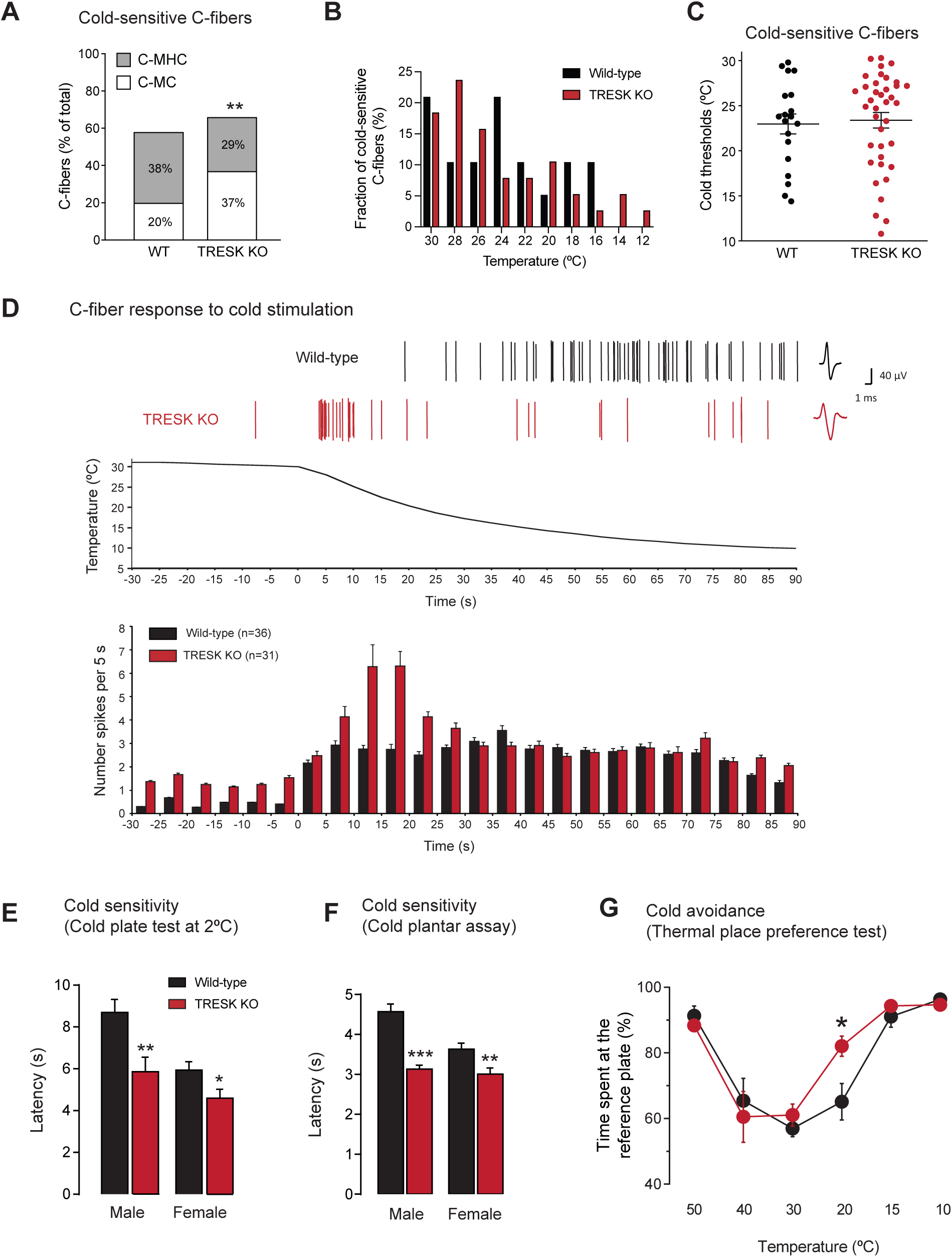
Cold allodynia in TRESK deleted mice. **A.** Fractions of cold-sensitive C-fibers in WT (total number of C-fibers = 35) and TRESK KO (n=37 C-fibers), measured in skin-nerve experiments. Numbers in bars are the percentage of Mechano-Heat-Cold (C-MHC) and Mechano-Cold (C-MC) fibers. A significant difference in the distribution of cold fibers is shown (**p<0.01, Chi-square test). **B.** Distribution of the C-fibers activated by cold at different temperatures between WT and KO animals. **C.** Distribution of cold thresholds from fibers recorded (WT n=19; KO n=38). Mean and SEM are shown. **D.** Representative experiments of C-fibers activated by a cold ramp. *Top*. Action potentials (Spikes) fired in response to a temperature decrease are shown for wild-type and a TRESK knockout cold C-fibers. The average action potential is presented on the right. A representative cold ramp from 30 to 10°C is shown below. *Bottom*: Histogram of mean responses to cooling, 5-second bin, of C-fibers from wild-type and TRESK KO mice. **E, F.** Cold sensitivity measured with the cold plate (2°C) and cold plantar assays in male and female wild-type and TRESK knockout animals (n=11-19 animals per group). **G.** Cold avoidance measured in the thermal place preference test. The percentage of time spent at the reference plate (30°C) at each experimental temperature is shown (n=8-14 male animals per group). Statistical differences are shown as *p<0.05, **p<0.01, ***p<0.001 (Student’s unpaired t-test) between WT and KO mice.

In good agreement with observations in C-fibers, behavioral responses to cold were also altered in animals lacking TRESK. Sensitivity to noxious cold (2°C; cold plate test) was enhanced in knockout animals from both sexes, with shorter latency times to elicit a nocifensive behavior (p=0.006 males; p=0.041 females, Fig 5E). In addition, cold sensitivity to more moderate temperatures assessed with the cold plantar assay was significantly higher in TRESK KO mice (p<0.0001 males, p=0.008 females, Fig 5F). Finally, we assessed whether changes in cold sensitivity modified the aptitude of mice to discriminate cool and cold temperatures in a test of thermal preference between plates at different temperatures. TRESK KO animals spent more time on the reference plate (at 30°C) than on the experimental plate at 20°C (p=0.022, Fig 5G) compared to wild-type animals, further supporting previous evidence from cold sensitive C-fibers recordings and cold sensitivity assays. In contrast, place preference assays at other temperatures (10, 15, 30, 40, 50°C) did not show significant alterations between mice genotypes, corroborating the non-involvement of TRESK in heat and warmth perception (Fig 4).

### Mice lacking TRESK show selective changes in persistent inflammatory and neuropathic pain

To assess the contribution of TRESK in chronic pain conditions, the behavior of TRESK KO and WT mice was measured after CFA injection into the mice hind paw, a model of persistent inflammatory pain. Both genotypes developed mechanical and thermal hypersensitivity in the injected hind paw beginning 1h after injection and lasting up to 16 days (Fig 6A). Despite the initial difference in basal mechanical thresholds (p=0.047), the extent of mechanical hyperalgesia in response to tactile stimuli was similar between WT and KO animals 1h after CFA injection and during the entire 16 days observation period. The development of thermal hyperalgesia was similar between both genotypes, thus showing that TRESK does not contribute significantly to peripheral sensitization of nociceptors due to inflammation. The sciatic nerve cuff-model was used to evaluate TRESK contribution in persistent neuropathic pain. Mechanical hypersensitivity developed in KO animals 5 and 7 days after sciatic nerve cuffing was significantly higher compared to WT (p=0.046 and p=0.047, respectively; Fig 6B). Sham surgery did not show significant effects on this parameter in any genotype (data not shown). Nevertheless, when the percentage of threshold decrease (difference vs. basal value) was compared between WT and KO animals, this showed similar values (at 5 days, KO: - 45.1±17.4%; WT: −37.9±12.1%; p=0.742; at 7 days, KO: −47.3±16.9%; WT: - 45.0±19.3%; p=0.931), indicating that nerve injury exerted a similar effect in both genotypes but, since KO had a lower mechanical threshold to begin with, animals reached lower mechanical thresholds at days 5 and 7. Mechanical hypersensitivity was undistinguishable between groups at later stages, at 14 and 21days post-injury. Heat sensitivity showed a similar level of hyperalgesia after neuropathy, akin to the lack of implication of TRESK in mechanical perception. Surprisingly withdrawal latencies from KO animals recovered baseline values faster than WT group after 14 and 21 days. Finally, we evaluated chemo-induced neuropathic pain with the anti-cancer drug oxaliplatin that induces cold allodynia in a majority of patients under therapy, to evaluate whether the development of cold hyperalgesia after oxaliplatin treatment was modified in TRESK KO. As expected, wild-type mice showed an enhanced cold sensitivity 90h after the oxaliplatin injection. Both, cold plantar assay (p=0.0002) and thermal place preference test (30/20°C; p=0.0045) showed significant differences compared to pre-injection values, indicating a higher sensitivity to cold stimuli in neuropathic mice (Fig 6C). In contrast, when TRESK KO animals were tested in the cold plantar assay, we did not observe any further decrease in paw withdrawal latency compared to the baseline value (p=0.912), which was already lower than that of WT animals. Again, thermal place preference between 20 and 30°C did not show significant differences after oxaliplatin treatment (p=0.770), suggesting that the enhanced cold sensitivity due to TRESK removal cannot be further increased by neuropathy induced by oxaliplatin.

**Figure 6:**
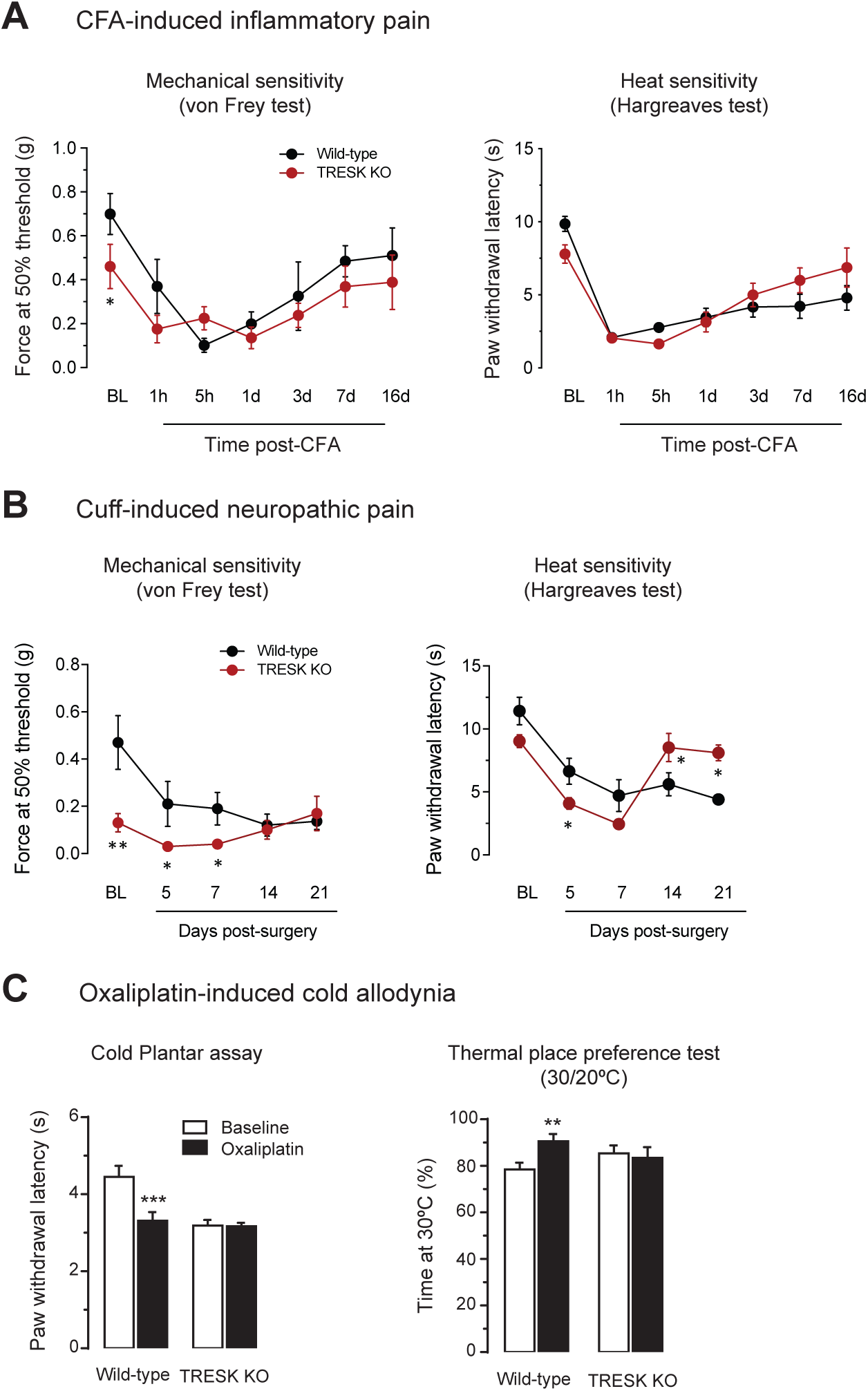
Changes in chronic pain in TRESK-deleted mice. **A.** Mechanical and thermal sensitivity in the CFA-induced inflammatory pain model. Mechanical sensitivity was measured with von Frey filaments (up and down method) and thermal sensitivity was measured with the Hargreaves test (n=7 animals in each group) **B.** Mechanical and thermal sensitivity in the cuff-induced neuropathic pain model. Mechanical sensitivity was measured with von Frey filaments (up and down method) and thermal sensitivity was measured with the Hargreaves test (n=6 animals in each group). **C.** Oxaliplatin-induced cold sensitization model. *Left.* Paw withdrawal latency to the cold plantar assay before (baseline) and 90h after oxaliplatin injection in WT and TRESK KO animals (n=9 and 10 animals per group). *Right.* Cold avoidance measured in the thermal place preference test. The percentage of time spent at the reference plate (30°C) versus the experimental temperature (20°C) is shown (n=6 and 8 animals per group). Statistical differences between WT and KO mice are shown as *p<0.05, **p<0.01, ***p<0.001 (Student’s unpaired t-test).

## Discussion

TRESK is a leak K^+^ channel highly expressed in human and rodent sensory neurons both in DRGs and TG [3-5,12,13,33-35]. Recent single-cell sequencing data indicates that in DRG and TG neurons, TRESK expression is restricted to some subtypes of sensory neurons, mainly, non-peptidergic medium/small diameter neurons involved in nociception [15,35-37]. In addition, TRESK is also present in a population of low-threshold mechanoreceptors (expressing high levels of TRKB and Piezo2) involved in touch sensation. TRESK main role has been attributed to preventing neuronal depolarization [1], and reduction of its expression after nerve injury or inflammation contribute to neuronal hyperexcitability [3,4,10]. Mutations in this channel have been linked to the enhanced nociceptor excitability that occurs during familial migraine with aura [20,21]. In the present study, we propose that TRESK balances the effects of depolarizing stimuli, acting as a brake to prevent the activation of specific subpopulations of sensory neurons expressing the channel. We found that TRESK channel prevents mechanical and cold hypersensitivity in mice, while the sensitivity to other stimulus modalities remains unaffected.

Small diameter nociceptive sensory neurons lacking TRESK presented a significant diminution in the amplitude of K^+^ current, an increase in membrane input resistance, and an increase in action potential duration, which are evidence of a major contribution of TRESK channel to K^+^ current in small diameter, presumably nociceptive, DRG neurons, in agreement with previous reports [5]. Resting membrane potentials were not different between TRESK KO and WT DRG neurons, which is in agreement with previous studies [3]. In contrast to other channels [11], it is likely that TRESK does not contribute significantly in setting this parameter but it has a major impact over the range of membrane potentials between the resting membrane potential and the action potential threshold. Indeed, other K^+^ channels from K_2P_ and other K^+^ channel families have been proposed to control membrane polarization at rest [38]. Although we cannot completely discard a compensatory effect due to other channels not analyzed (e.g. HCN, KCNQ), TRESK removal does not seem to significantly modify the expression of other leak channels expressed in sensory neurons that have a major contribution to the K^+^ background current (TREK-1, TREK-2, TRAAK) [5]. Therefore, the effects observed in excitability appear to be mainly attributable to the specific removal of TRESK. Nociceptive DRG neurons from TRESK KO mice showed higher excitability, with a decreased injected current threshold for action potential firing and an increased firing response to a depolarizing ramp, which is also consistent with a significant decrease of a potassium conductance in KO mice. Despite finding significant differences in these parameters, these effects could be underestimated by the fact that TRESK is not expressed in all types of nociceptive sensory neurons. Therefore, it is possible that studying specific subpopulations of sensory neurons (mainly non-peptidergic) genetically labeled with specific markers might render larger effects in excitability after TRESK deletion. Studies in this line are underway to unveil particular effects in different types of sensory neurons. Similar findings on neuronal excitability to these found in the present study have been reported after a decreased TRESK expression, such as in a functional TRESK[G339R] knockout mice [3,23], after sciatic nerve axotomy [4] or in a model of cancer-associated pain [24]. This is in agreement with the neuronal activation observed by compounds blocking TRESK [4,33,39]. Conversely, opposite consequences on excitability were found when overexpressing TRESK channel [18,24], thus indicating that the regulation of TRESK expression is an important factor to control sensory neuron excitability.

Recordings from C-fibers showed that detection of mechanical and cold stimuli, but no heat, were enhanced in the KO mice. The effects of TRESK KO were specific to some properties of these modalities and did not enhance neurons excitability over the whole range of stimulation. This indicates that removal of TRESK channel does not produce a general increase in excitability but a selective effect on certain types of fibers/sensory neuron subtypes affecting specific sensory modalities. This is similar to what occurs for other K_2P_ or K^+^ channels: genetic deletion of TREK-1 modifies mechanical, heat and cold pain perception [7] but when TRAAK is deleted, cold sensitivity remains unaffected and animals only present an enhanced heat sensitivity. Interestingly, deleting both channels (TREK-1/TRAAK double KO mice) potentiates the effects on cold sensitivity compared to TREK-1 alone [9]. Besides, the role of TREK-2 in thermosensation is different from that of TREK-1 and TRAAK channels and deletion of the channel enhances thermal sensitivity to non-aversive warm temperatures but not to cold stimuli [8]. Besides, knocking down the BK_Ca_ potassium channel does not modify acute nociceptive or neuropathic pain, but KO mice show an increased nociceptive behavior in models of persistent inflammatory pain [40], highlighting its specific participation during inflammation-induced hyperalgesia. Our data shows that the lack of TRESK enhances cold sensitivity by increasing the percentage of C-fibers activated at moderate cold temperatures, which is translated at the behavioral level by an enhanced sensitivity to cold stimuli. The bigger fraction of cold sensitive C-MC fibers (37%) in KO animals compared to wild-types (20%) suggests that TRESK normally silences a population of neurons that would be only activated at low temperatures in normal conditions. This is combined with a decrease in the fractions of C-MHC and C-MH fibers. These observations are consistent with a possible role of TRESK in preventing cold allodynia, acting as a brake to avoid C-fiber activation at moderate cool temperatures. In fact, knocking out TRESK mimics cold allodynia and oxaliplatin injection is not able to further enhance cold sensitivity. Cold allodynia induced by the anti-cancer drug oxaliplatin has been attributed to a combined remodeling of ion channels in subsets of sensory neurons, including down-regulation of TREK-1, TREK-2, TRAAK, K_v_1.1 and K_v_1.2, coupled with up-regulation of TRPA1, Na_v_1.8 and HCN1, while other channels involved, such as TRPM8, are not significantly modified [8,41]. It has been proposed that a reduced activity of K_2P_ channels when temperature decreases is responsible of releasing the excitability brake exerted by these channels, and thus, combined with depolarizing thermo-TRP channels activation, trigger sensory neurons activity when they sense cold temperature stimuli [5,8,9]. TRESK activity is not significantly modified by temperature between 20-37°C [5]. Since we show that DRG neurons from TRESK KO have lower action potential threshold and increase firing upon membrane depolarization, it is possible that the channel prevents the activation of a subpopulation of C-fibers by cold-activated excitatory depolarizing channels even if lower temperature does not modify its activity, at least, in a moderate range. Moreover, we show an increase in input membrane resistance in TRESK KO DRG neurons. Low temperatures increase the membrane resistance, which potentiates voltage change by membrane currents and enhance excitability. This effect has been shown previously to contribute to cold sensation [42].

TRPA1 was initially proposed to be involved in noxious cold sensing [43] but recent reports have questioned its role. It has been hypothesized that it might be regulating cold sensitivity indirectly, rather than acting as a cold-sensing receptor [32,44]. We observed a reduced response to TRPA1 agonists (Fig 3) but no change in its mRNA expression in DRGs. Whether the enhanced cold sensitivity in TRESK KO animals being dependent on TRPA1 expression remains a possibility, recent studies have shown that TRPA1-expressing neurons are unresponsive to noxious cold stimuli [44]. K_v_ channels can blunt cold responses by opposing to depolarizing stimuli in TRPA1^+^ and TRPM8^+^ neurons [45,46]. It is possible that TRESK might exert a similar effect, hence silencing cold responses in normal conditions. Neurons lacking TRESK or down-regulation of the channel might release this excitability brake, uncovering an enhanced cold sensation. Even if this is the case, it remains to be examined in greater depth. Moreover, changes observed in the sensitivity to AITC might result from post-transcriptional regulation of protein expression levels or other intracellular signaling mechanisms altered in TRESK KO cellular background such as phosphorylation of the receptor due to the enhanced excitability of TRESK KO neurons.

In addition to cold, mechanical sensitivity is also enhanced in TRESK KO animals, which present a higher number of C-fibers with low mechanical thresholds (<12 mN) and an increased response to von Frey hairs application. Similar results were previously reported after knocking down TRESK expression or after injecting TRESK blockers, where animals showed mechanical allodynia [4,24,39]. Likewise, trigeminal expression of a mutated form of TRESK linked to migraine results in facial mechanical allodynia [21]. Similar to cold, mechanical sensitivity is enhanced by decreasing the threshold of activation of a population of sensory fibers, and therefore, a higher number of fibers are activated by low intensity mechanical stimulation. This effect was not linked to gender, since both male and female animals showed similar responses. Piezo2 channel is involved in the detection of both innocuous touch and high threshold noxious mechanical stimuli due to its expression in low threshold mechanoreceptors (LTMR) and nociceptors [47-50], and has been proposed to play a crucial role in the generation of mechanical allodynia during inflammation and neuropathic pain states [47]. TRESK is also highly expressed in these populations of sensory neurons involved in mechanical hypersensitivity [15]. We can hypothesize that when these neuronal populations lack TRESK, they are more sensitive to mechanical stimuli activating Piezo2 or other mechanosensitive channels such as the recently identified TACAN [51], that would depolarize sensory neurons more effectively. We have not evaluated whether LTMR lacking TRESK are more excitable and if other touch modalities such as light touch (brush stroke) are also enhanced by TRESK deletion, but mechanical allodynia present in these animals suggests that they might also be affected. TRESK is notably expressed in MrgprD^+^ neurons [15] and deletion of this subpopulation of neurons abolishes mechanical but not thermal pain [52]. Here, the opposite effect is found after TRESK deletion and that will likely enhance the excitability of MrgprD^+^ neurons. In contrast, mechanical sensitization that occurs after inflammation does not seem to be further enhanced by the absence of TRESK whereas mechanical hypersensitivity after nerve injury seems to reach lower values compared to wild-type animals. Nevertheless, this effect appears to be related to the lower mechanical threshold present in TRESK KO animals since the mechanical threshold decrease is similar in both WT and KO groups. Our data suggests that the molecular effect of injury, down-regulating TRESK expression [4], has already been achieved by genetic removal of TRESK, thus any further decrease in mechanical threshold observed is likely due to other mechanisms sensitizing nociceptors or derived from central effects.

In our study, TRESK KO mice did not display relevant differences in heat sensitivity to the tested temperatures. Only a significant difference on the thermal place preference test was observed between 20/30°C that seems to reflect the observed enhancement of cold sensitivity rather than any effect on the warm/heat sensitivity. In agreement, TRESK KO C-fibers recorded did not show any specific effects upon temperature increase compared to WT. No significant alterations on heat sensitivity were either observed after CFA-induced inflammation, and only a slightly higher effect on heat sensitivity was observed in the neuropathic pain model of mice lacking TRESK. All together it seems that the role of TRESK in neurons detecting warm or hot stimuli is not relevant, in contrast to other channels of the TRP and K_2P_ family that play a major role [7-9,44,53,54]. Chemical sensitivity was altered for some stimuli (AITC, osmosensitivity) but not for others (capsaicin, menthol). Besides the altered sensitivity to AITC as previously discussed, removal of TRESK produced a decreased response to pain induced by osmotic stimuli. Whether TRESK is involved in detecting this type of stimuli is unknown, although we previously described that hypertonic and hypotonic stimuli are able to modify TRESK currents [55]. Interestingly, similar results in osmotic pain have been reported in animals lacking channels of the TREK subfamily [7-9] and in TRPV4 KO animals [29]. Similarly to TREK-1/2 KO mice but not TRAAK KO, PGE_2_ was unable to sensitize nociceptors to osmotic stimuli in TRESK KO animals. It has been proposed that the negative regulation of TREK-1/2 channels by cAMP/PKA downstream of the G-coupled PGE_2_ receptor might be involved in the lack of PGE_2_-mediated nociceptor sensitization. PKA is known to phosphorylate TRESK and keep it in a resting state where the channel is less active but a direct link between PKA activation, TRESK activity and osmotic sensing is still unknown. In fact, neurons activated by radial stretch and hypotonic stimuli seem to express TRESK, as they are sensitive to hydroxy-α-sanshool, a TRESK blocker [56]. These neurons were initially classified as LTMR or non-peptidergic nociceptors, which is in agreement with the TRESK expression pattern obtained from single-cell mRNA sequencing [15].

In summary, we describe that genetic removal of TRESK enhances mechanical and cold sensitivity in mice; consequently TRESK function is important to regulate the excitability in specific subpopulations of sensory neurons involved in mechanosensitivity and cold sensing. Because mice lacking TRESK display mechanical hypersensitivity as well as enhanced cold perception similar to the reported cold allodynia induced by oxaliplatin treatment, activators of the channel would be expected to improve these conditions during neuropathic pain. The development of specific channel openers/activators would be worthwhile and due to the rather restricted expression of TRESK in sensory ganglia (DRG and TG) will avoid undesired side effects in other tissues.

## Methods

### Animals

All behavioral and experimental procedures were carried out in accordance with the recommendations of the International Association for the Study of Pain (IASP) and were reviewed and approved by the Animal Care Committee of the University of Barcelona and by the Department of the Environment of the *Generalitat de Catalunya*, Catalonia, Spain (#8466, 8468, 8548, 6869, 9876). Female and male C57BL/6N mice between 8 and 15 weeks old were used in all experimental procedures (RNA extraction and qPCR, cell cultures, electrophysiology, calcium imaging, nerve fiber recording and behavior) unless differentially indicated. Mice were housed at 22°C with free access to food and water in an alternating 12h light and dark cycle.

TRESK (Kcnk18/K2P18.1) knockout mice and wild-type (WT) littermates were obtained from the KOMP Repository (Mouse Biology Program, University of California, Davis, CA). The TRESK knockout (KO) mouse was generated by replacing the complete Kcnk18 gene by a ZEN-UB1 cassette according to the VelociGene’s KOMP Definitive Null Allele Design. At 3 weeks of age, WT or KO newborn mice were weaned, separated, and identified by ear punching. Genomic DNA was isolated from tail snip samples with Maxwell® Mouse Tail DNA Purification Kit (Promega, Madison, WI). Polymerase chain reaction (PCR) was performed with primers to detect the Kcnk18 gene: forward 5’-ACCAACACCAAGCTGTCTTGTTTCTC-3’ and reverse 5’-AGACAGATGGACGGACAGACATAGATG-3’ or the inserted cassette in the KO mice: forward (REG-Neo-F) 5’-GCAGCCTCTGTTCCACATACACTTCA-3’ and reverse (gene-specific) 5’-AGACTTCTCCCAGGTAACAACTCTGC-3’. The PCR mixture contained 1 μl DNA sample, 2.5 μl PCR buffer (10x concentration), 2 μl dNTP mixture (2.5 mM), 0.5 μl (20 μM) forward and reverse primers, 0.2 μl Taq DNA polymerase (5 U/μl), 1.7 μl MgCl_2_ (25 mM), 6.5 μl Betaine (5 M), 0.325 μl DMSO and 9.7 μl water (final volume of 25 μl). PCR amplifications were carried out with 31 cycles in a programmable thermal cycler (Eppendorf AG, Hamburg, Germany). The program used was: 94°C for 5 min and cycles of 94°C for 15 s, 60°C for 30 s, 72°C for 40 s, with a final extension at 72°C for 5 min. PCR products were analyzed by electrophoresis in 1% agarose gels. Once identified, genotyped animals were used as breeders for colony expansion and their offspring were used in all experimental procedures in which mice were required.

### Behavioral studies

Female and male wild-type or TRESK KO mice between 8 and 15 weeks of age were used in all behavioral studies. To avoid stress-induced variability in the results, mice were habituated to the experimental room and the experimental setup prior testing. Behavioral measurements were done in a quiet room, taking great care to minimize or avoid discomfort of the animals.

### Mouse mechanical sensitivity

Mechanical sensitivity of wild-type and TRESK KO mice was assessed using the ‘up and down’ method by the application of calibrated von Frey filaments (North Coast Medical, Inc. Morgan Hill, CA) as previously described [4,39]. The von Frey filaments [size: 2.44, 2.83, 3.22, 3.61, 3.84, 4.08, 4.17, 4.31, 4.56; equivalent to (in grams) 0.04, 0.07, 0.16, 0.4, 0.6, 1, 1.4, 2 and 4] were applied perpendicularly to the plantar surface of the hind paw and gently pushed to the bending point for 5 s. The 50% withdrawal threshold was determined using the up and down method [57]. A brisk hind paw lift in response to von Frey filament stimulation was regarded as a withdrawal response. Dynamic plantar aesthesiometer (Ugo Basile, Italy) was also used to assess mechanical sensitivity. A von Frey-type 0.5 mm filament was applied with a 10 s ramp (0 to 7.5 g) and the hind paw withdrawal threshold of mice was recorded.

### Mouse thermal sensitivity: Hot plate and cold plate test

After habituation, the cold/hot plate apparatus (Ugo Basile, Italy) was set to 2°C, 52°C or 56°C and animals were individually placed in the center of the plate. Latency time to elicit a nocifensive behavior (a jump or a paw lick/lift) was counted with the apparatus stopwatch and the average of three separately trials was used as a measurement. Since the temperatures tested are in the noxious range, a cut-off time of 25 seconds was established to avoid tissue damage.

### Dynamic hot plate test

In contrast to the conventional hot plate, the dynamic hot plate allows the testing of a wide range of temperatures. Before testing, animals were habituated and later placed individually to the hot/cold plate apparatus where the plate temperature increased from 30°C to 50°C at a 1°C/min rate. To determine the temperature that is perceived as noxious for mice and quantify pain-related behaviors, the number of jumps at each temperature was scored.

### Radiant heat test

The heat sensitivity of mice was assessed by measuring hind paw withdrawal latency from a radiant infrared source (Hargreaves’ method) using the Ugo Basile (Italy) Model 37370 Plantar test. Each measurement was the mean of 3 trials spaced 15 min apart. For all experiments, infrared intensity was set to 30% and a cut-off time of 20 seconds was established to avoid skin burn damage.

### Thermal place preference test

The thermal place preference test is a test of better comfort temperature rather than an indicator of temperature aversion. The hot/cold plate apparatus was placed side by side with a complementary plate and a small divider platform was situated between them to connect the two devices (Ugo Basile, Italy). The reference plate was always set at 30°C and the test plate was set at 50, 40, 30, 20, 15 or 10°C. Animals were habituated at the experience room for 30 minutes before testing and then they were allowed to investigate the testing setup for 10 minutes or until animals crossed homogeneously from one plate to the other. Mice were then placed individually to the center of the platform and once they crossed to a plate the cumulative time they spent on each plate for a total of 5 minutes was then counted. Only animals that performed properly when both plates were set to 30°C were used for the study (50% of the time at each plate and more than 2 crossings). To exclude the possibility that animals could be learning which one the reference plate was, the position of the reference and the test plate was switched randomly between trials.

### Cold plantar assay

To complement the cold plate test, we studied noxious cold sensitivity of TRESK KO mice using the cold plantar assay [58]. This assay produces an unambiguous nocifensive response that is easily identified when compared to the cold plate test. Animals were placed on top of a 1/8’’ thick glass plate and were enclosed in transparent boxes separated by opaque dividers to prevent animals to see each other. The cold probe consisted of a modified 3 mL syringe filled with freshly powdered dry ice. This powder was then packed into a pellet and its surface was flattened. Using a mirror to target the mouse hind paw, we applied the dry ice pellet below the glass, making sure that the paw was completely in contact with it. This delivers a cooling ramp to the mice paw and a few seconds later withdrawal responses occur. The withdrawal latency time was measured with a stopwatch and the final withdrawal latency time for each animal was the average of 3 trials, which were tested at intervals of at least 15 minutes. A cut-off time of 20 seconds was used to prevent tissue damage.

### Evaluation of nocifensive behavior

Using a 30g needle, 10 μl of a solution containing 100 μM AITC (10%), capsaicin (1μg/10μl) or their vehicle solutions were administered intradermally into the plantar surface of the hind paw. Mice behavior was observed and the number of flinches and lickings of the paw were counted for a 5-minute period starting immediately after the injection. On the day previous to testing, animals were habituated to the testing room and to the handling procedure. The flinching and licking test was also used to examine the painful response of mice to different osmolality solutions, both in naive and sensitized conditions. 10 μl of the following solutions were administered in the hind paw to different groups of male mice: NaCl −33% (hypotonic, 100 mOsm·Kg^-1^), NaCl 2% (hypertonic, 622 mOsm·Kg^-1^), NaCl 10% (hypertonic, 3157 mOsm·Kg^-1^) and PBS (Isotonic, 298 mOsm·Kg^-1^). A different group of animals was used to study osmotic pain under inflammatory conditions. A 5 μl injection of prostaglandin E_2_ (PGE_2_; 10 μM) was injected into the hind paw of each mouse and 30 minutes after, 10 μl of NaCl - 33% or NaCl 2% were injected into the sensitized paw. After the injection of different osmolality solutions, nocifensive behaviors of paw licking and shaking were manually counted for 5 minutes.

### CFA model of inflammatory pain

After baseline measurements and under brief isoflurane anesthesia, Complete Freund’s Adjuvant (CFA, Sigma-Aldrich; 20 μl; 1mg/ml) was injected subcutaneously (glabrous skin) in the hind paw of mice to induce a local inflammation. Mechanical von Frey threshold and heat withdrawal latency (radiant heat test) was examined at different time points after the injection (1h, 5h, 1d, 3d, 7d and 16d).

### Formalin test

After habituation, mouse hind paw was subcutaneously injected with formalin (10 μl of a 5% formaldehyde solution) and nocifensive behaviors were measured for 50 minutes after the injection. The cumulative time spent shaking and licking the injected paw was counted with a stopwatch in 5 min periods.

### Oxaliplatin-induced cold hypersensitivity

Prior to oxaliplatin injection, naive thermal preference was measured with the thermal place preference test set at 20°C vs. 30°C. Thirty min later, paw withdrawal latency to cold stimuli was determined using the cold plantar assay. A single intraperiotoneal injection of oxaliplatin (6 mg/Kg in PBS with 5% glucose) was delivered to mice. Cold preference and cold sensitivity were re-evaluated 90 h after oxaliplatin injection, a time point that is known to correlate with the peak of oxaliplatin-induced cold hyperalgesia [41].

### Cuff-induced neuropathic pain model

The sciatic nerve cuffing model was used to induce mechanical and thermal hyperalgesia in the hind paw of mice, as described in [59]. Mice were housed individually to avoid stress derived from the surgery and to prevent injury. Cardboard rolls were placed into their home cages to provide shelter and additional stimulation after the surgery. All surgeries were done under aseptic conditions using intraperitoneal ketamine/xylazyne anesthesia. The left leg of mice was shaved from the hip to the knee and the surgical field was disinfected. A 0.5 cm incision parallel to the femur was made to expose the common branch of the sciatic nerve, which appeared after separating the muscles close to the femur. A drop of sterile physiological saline was applied to prevent the nerve from dehydrating and this concluded the procedure for the sham group. The ‘cuff’ consisted of a 2 mm-long piece of polyethylene tubing (PE-20) with an inner diameter of 0.38 mm and an outer diameter of 1.09 mm, opened by one of its sides. For the cuff group, the sciatic nerve was gently straightened with two sterile sticks and the ‘cuff’ was inserted around the main branch of the nerve and closed by applying moderate pressure with surgical forceps. To ensure that the ‘cuff’ was correctly positioned and closed, it was turned gently around the nerve. Both sham and cuff surgeries ended by suturing the incision with surgical knots. After the procedure, animals were placed in their respective home cages laying on their right side and were under constant supervision until they were completely awakened. Mechanical and heat sensitivity of mice were determined with the von Frey and the radiant heat tests as previously described, before surgery and 5, 7, 14 and 21 days after surgery.

### Skin-nerve preparation and single fiber recordings

The isolated skin-saphenous nerve preparation for single C-fiber recording was used as previously described [9]. The hind paw skin of male mice 10 to 20 weeks of age was isolated with the saphenous nerve. The skin was pinned corium side up in a perfusion chamber with the nerve being pulled in a recording chamber filled with paraffin oil. The skin was perfused with warm (∼30-31°C) synthetic interstitial fluid (SIF), in mM: 120 NaCl, 3.5 KCl, 5 NaHCO_3_, 1.7 NaH_2_PO_4_, 2 CaCl_2_, 0.7 MgSO_4_, 9.5 Na-Gluconate, 5.5 glucose, 7.5 sucrose, and 10 HEPES, pH 7.4 adjusted with NaOH, saturated with O_2_/CO_2_ 95%/5%. Isolated nerve fibers were placed on a gold recording electrode connected to a DAM-80 AC differential amplifier (WPI), Digidata 1322A (Axon Instruments) and Spike2 software (CED) to record extracellular potentials. The skin was probed with mechanical stimulation with a glass rod and calibrated von Frey filaments to characterize the mechanical sensitivity of nerve fibers. The C-fibers were classified according to their conduction velocity of less than 1.2 m/s measured by electrical stimulation (Stimulus Isolator A385, WPI). The skin receptive field of C-fibers was isolated from the surrounding bath chamber with a stainless steel ring 0.8 cm in diameter, an internal volume of 400 μl, and a hot or cold temperature-controlled SIF was infused into the ring through a bipolar temperature controller CL-100 (Warner instrument). Electrical recordings were amplified (x10 000), band-pass filtered between 10 Hz and 10 kHz and stored on computer at 20 kHz. The action potentials were detected and analyzed offline with the principal component analysis extension of the Spike2 software (CED).

### RNA extraction and Quantitative real-time PCR

Mouse tissue samples were obtained from dorsal root ganglia, kept in RNAlater solution (Ambion) and stored at −80°C until use. Total RNA was isolated using the Nucleospin RNA (Macherey-Nagel) and first-strand cDNA was then transcribed using the SuperScript IV Reverse Transcriptase (Invitrogen, ThermoFisher Scientific) according to the manufacturer’s instructions. Quantitative real-time PCR was performed in an ABI Prism 7300 using the Fast SYBR Green Master mix (Applied Biosystems) and primers detailed in Suppl. Table 1; obtained from Invitrogen (ThermoFisher Scientific). Amplification of Glyceraldehyde 3-phosphate dehydrogenase (GAPDH) transcripts was used as a standard for normalization of all qPCR experiments and gene-fold expression was assessed using the ΔC_T_ method. All reactions were performed in triplicate. After amplification, melting curves were obtained and evaluated to confirm correct transcript amplification.

### Culture of dorsal root ganglion neurons

Mice were euthanized by decapitation under anesthesia (isoflurane) and thoracic, lumbar and cervical dorsal root ganglia (DRG) were removed for neuronal culture as previously described [4,55]. Briefly, DRGs were collected and maintained in cold (4–5°C) Ca^2+^ - and Mg^2+^-free Phosphate Buffered Saline solution (PBS) supplemented with 10 mM glucose, 10 mM Hepes, 100 U.I./mL penicillin and 100 μg/mL streptomycin until dissociation. Subsequently, ganglia were incubated in 2 ml HAM F-12 with collagenase CLS I (1 mg/ml; Biochrome AG, Berlin) and Bovine Serum Albumin (BSA, 1 mg/ml) for 1 h 45 min at 37°C followed by 15 min trypsin treatment (0.25%). Ganglia were then resuspended in Dulbecco’s Modified Eagle medium (DMEM) supplemented with 10% FBS, penicillin/streptomycin (100 μg/ml) and L-glutamine (100 mg/ml) and mechanical dissociation was conducted with fire-polished glass Pasteur pipettes of decreasing diameters. Neurons were centrifuged at 1000 rpm for 5 min and re-suspended in culture medium [DMEM + 10% FBS, 100 μg/ml penicillin/streptomycin, 100 mg/mL L-glutamine]. Cell suspensions were transferred to 12 mm-diameter glass coverslips pre-treated with poly-L-lysine/laminin and incubated at 37°C in humidified 5% CO_2_ atmosphere for up to 1 day, before being used for patch-clamp electrophysiological recordings or calcium imaging experiments. Nerve Growth Factor or other growth factors were not added.

### Calcium imaging

Cultured DRG neurons from wild-type and TRESK KO mice were loaded with 5 μM fura-2/AM (Invitrogen, Carlsbad, CA) for 45-60 min at 37°C in culture medium. Coverslips with fura-2 loaded cells were transferred into an open flow chamber (0.5 ml) mounted on the stage of an inverted Olympus IX70 microscope equipped with a TILL monochromator as a source of illumination. Pictures were acquired with an attached cooled CCD camera (Orca II-ER, Hamamatsu Photonics, Japan) and stored and analyzed on a PC computer using Aquacosmos software (Hamamatsu Photonics, Shizuoka, Japan). After a stabilization period, pairs of images were obtained every 4 s at excitation wavelengths of 340 (λ1) or 380 nm (λ2; 10 nm bandwidth filters) in order to excite the Ca^2+^ bound or Ca^2+^ free forms of the fura-2 dye, respectively. The emission wavelength was 510 nm (12-nm bandwidth filter). Typically, 20-40 cells were present in the microscope field. [Ca^2+^]_i_ values were calculated and analyzed individually for each single cell from the 340- to 380-nm fluorescence ratios at each time point. Only neurons that produced a response >10% of the baseline value and that, at the end of the experiment, produced a Ca^2+^ response to KCl-induced depolarization (50 mM) were included in the analysis. Several experiments with cells from different primary cultures and different animals were used in all the groups assayed. The extracellular (bath) solution used was 140 mM NaCl, 4.3 mM KCl, 1.3 mM CaCl_2_, 1 mM MgCl_2_, 10 mM glucose, 10 mM HEPES, at pH 7.4 with NaOH. Experiments were performed at room temperature.

### Electrophysiological recording

Electrophysiological recordings in DRG sensory neurons were performed as previously described [39,55,60]. Briefly, recordings were performed with a patch-clamp amplifier (Axopatch 200B, Molecular Devices, Union City, CA) and restricted to small size DRG neurons (<30 μm soma diameter), which largely correspond to nociceptive neurons [61]. Patch electrodes were fabricated in a Flaming/Brown micropipette puller P-97 (Sutter instruments, Novato, CA). Electrodes had a resistance between 2-4 MΩ when filled with intracellular solution (in mM): 140 KCl, 2.1 CaCl_2_, 2.5 MgCl_2_, 5 EGTA, 10 HEPES, 2 ATP at pH 7.3. Bath solution (in mM): 145 NaCl, 5 KCl, 2 CaCl_2_, 2 MgCl_2_, 10 HEPES, 5 glucose at pH 7.4. The osmolality of the isotonic solution was 310.6±1.8 mOsm/Kg. Membrane currents were recorded in the whole-cell patch-clamp configuration, filtered at 2 kHz, digitized at 10 kHz and acquired with pClamp 10 software. Data was analyzed with Clampfit 10 (Molecular Devices) and Prism 7 (GraphPad Software, Inc., La Jolla, CA). Series resistance was always kept below 15 MΩ and compensated at 70-80%. All recordings were done at room temperature (22-23°C), 18-24h after dissociation. To study sensory neuron excitability, after achieving the whole-cell configuration in the patch clamp technique, the amplifier was switched to current-clamp bridge mode. Only neurons with a resting membrane voltage below −50 mV were considered for the study. To study neuronal excitability, we examined the resting membrane potential (RMP); action potential (AP) current threshold elicited by 400 ms depolarizing current pulses in 10 pA increments; whole-cell input resistance (R_in_) was calculated on the basis of the steady-state I-V relationship during a series of 400 ms hyperpolarizing currents delivered in steps of 10 pA from −50 to −10 pA; AP amplitude was measured from RMP to AP peak and AP duration was measured at 50% of AP amplitude.

### Drugs

All reagents and culture media were obtained from Sigma-Aldrich (Madrid, Spain) unless otherwise indicated. Menthol (100 μM), allyl isothiocyanate (AITC; 100 μM) and capsaicin (1 μM) were also purchased from Sigma (Madrid, Spain).

### Data analysis

Data are presented as mean ± SEM. Statistical differences between different sets of data were assessed by performing paired or unpaired Student’s t-tests or Wilcoxon matched pairs test, Chi-square test or Fisher’s exact test, as indicated. The significance level was set at p<0.05 in all statistical analyses. Data analysis was performed using GraphPad Prism 7 software and GraphPad QuickCalcs online tools (GraphPad Software, Inc., La Jolla, CA)

## Acknowledgements

Supported by grants from Ministerio de Economia y Competitividad and Instituto de Salud Carlos III/FEDER of Spain FIS PI14/00141 (XG), FIS PI17/00296 (XG), RETICs Oftared RD16/0008/0014 (XG), Generalitat de Catalunya 2017SGR737 (XG) and Ministerio de Economia, Industria y Competitividad, Spain (BFU2017-83317-P (DS).

## Authors’ contributions

Authors AC, AAB and GC performed electrophysiological recordings in neurons and calcium imaging. AC, APC and AAB performed behavioral experiments. AC, AAB carried out primary cell cultures. AC, GC, APC and NC performed qPCR experiments. AN and JN performed skin-nerve recordings. AC, DS, JN, NC and XG participated in the design of the study and performed the statistical analysis. XG conceived the study, oversaw the research and prepared the manuscript with help from all others. All authors read and approved the final manuscript.

---

**Supplementary Figure 1: Dynamic hot plate** In contrast to the conventional hot plate, the dynamic hot plate allows the testing of a wide range of temperatures. Plate temperature was ramped from 30°C to 50°C at a 1°C/min rate. To determine the temperature that is perceived as noxious for mice and quantify pain-related behaviors, the number of jumps at each temperature was scored. *Left*: number of jumps elicited at each temperature in male (top) and female (bottom) in wild-type and TRESK KO mice (n=11-15 animals per group). Only a significant difference was obtained at 49°C (p<0.05, unpaired t-test) between wild-type and TRESK KO females. *Right, top*: mean temperature at which animals did the first jump (threshold). Right. bottom: Total number of jumps for male and female mice in the whole range of temperatures. Values for temperatures between 30 and 38°C are not shown in the plots since were probably detected as non-noxious and did not produce any observable response.

## References

1. Enyedi P, Czirják G. Molecular background of leak K+ currents: two-pore domain potassium channels. Physiological Reviews. 2010 Apr;90(2):559–605.

2. Yamamoto Y, Hatakeyama T, Taniguchi K. Immunohistochemical colocalization of TREK-1, TREK-2 and TRAAK with TRP channels in the trigeminal ganglion cells. Neurosci Lett. 2009 Apr 24;454(2):129–33.

3. Dobler T, Springauf A, Tovornik S, Weber M, Schmitt A, Sedlmeier R, Wischmeyer E, Döring F. TRESK two-pore-domain K+ channels constitute a significant component of background potassium currents in murine dorsal root ganglion neurones. J Physiol (Lond). 2007 Dec 15;585(Pt 3):867–79.

4. Tulleuda A, Cokic B, Callejo G, Saiani B, Serra J, Gasull X. TRESK channel contribution to nociceptive sensory neurons excitability: modulation by nerve injury. Mol Pain. 2011;7(1):30.

5. Kang D, Kim D. TREK-2 (K2P10.1) and TRESK (K2P18.1) are major background K+ channels in dorsal root ganglion neurons. Am J Physiol, Cell Physiol. 2006 Jul;291(1):C138–46.

6. Patel AJ, Honoré E, Maingret F, Lesage F, Fink M, Duprat F, Lazdunski M. A mammalian two pore domain mechano-gated S-like K+ channel. EMBO J. 1998 Aug 3;17(15):4283–90.

7. Alloui A, Zimmermann K, Mamet J, Duprat F, Noël J, Chemin J, Guy N, Blondeau N, Voilley N, Rubat-Coudert C, Borsotto M, Romey G, Heurteaux C, Reeh P, Eschalier A, Lazdunski M. TREK-1, a K+ channel involved in polymodal pain perception. EMBO J. 2006 Jun 7;25(11):2368–76.

8. Pereira V, Busserolles J, Christin M, Devilliers M, Poupon L, Legha W, Alloui A, Aissouni Y, Bourinet E, Lesage F, Eschalier A, Lazdunski M, Noel J. Role of the TREK2 potassium channel in cold and warm thermosensation and in pain perception. Pain. 2014 Dec;155(12):2534–44.

9. Noël J, Zimmermann K, Busserolles J, Deval E, Alloui A, Diochot S, Guy N, Borsotto M, Reeh P, Eschalier A, Lazdunski M. The mechano-activated K+ channels TRAAK and TREK-1 control both warm and cold perception. EMBO J. 2009 May 6;28(9):1308–18.

10. Marsh B, Acosta C, Djouhri L, Lawson SN. Leak K⁺ channel mRNAs in dorsal root ganglia: relation to inflammation and spontaneous pain behaviour. Molecular and cellular neurosciences. 2012 Mar;49(3):375–86.

11. Acosta C, Djouhri L, Watkins R, Berry C, Bromage K, Lawson SN. TREK2 Expressed Selectively in IB4-Binding C-Fiber Nociceptors Hyperpolarizes Their Membrane Potentials and Limits Spontaneous Pain. J Neurosci. Society for Neuroscience; 2014 Jan 22;34(4):1494–509.

12. Manteniotis S, Lehmann R, Flegel C, Vogel F, Hofreuter A, Schreiner BSP, Altmüller J, Becker C, Schöbel N, Hatt H, Gisselmann G. Comprehensive RNA-Seq Expression Analysis of Sensory Ganglia with a Focus on Ion Channels and GPCRs in Trigeminal Ganglia. Zhang Z, editor. PLoS ONE. Public Library of Science; 2013 Nov 8;8(11):e79523.

13. Ray P, Torck A, Quigley L, Wangzhou A, Neiman M, Rao C, Lam T, Kim J-Y, Kim TH, Zhang MQ, Dussor G, Price TJ. Comparative transcriptome profiling of the human and mouse dorsal root ganglia: an RNA-seq-based resource for pain and sensory neuroscience research. Pain. 2018 Jul;159(7):1325–45.

14. Flegel C, Schöbel N, Altmüller J, Becker C, Tannapfel A, Hatt H, Gisselmann G. RNA-Seq Analysis of Human Trigeminal and Dorsal Root Ganglia with a Focus on Chemoreceptors. PLoS ONE. 2015;10(6):e0128951.

15. Usoskin D, Furlan A, Islam S, Abdo H, Lönnerberg P, Lou D, Hjerling-Leffler J, Haeggström J, Kharchenko O, Kharchenko PV, Linnarsson S, Ernfors P. Unbiased classification of sensory neuron types by large-scale single-cell RNA sequencing. Nat Neurosci. 2015 Jan;18(1):145–53.

16. Zeisel A, Hochgerner H, Lönnerberg P, Johnsson A, Memic F, van der Zwan J, Häring M, Braun E, Borm LE, La Manno G, Codeluppi S, Furlan A, Lee K, Skene N, Harris KD, Hjerling-Leffler J, Arenas E, Ernfors P, Marklund U, Linnarsson S. Molecular Architecture of the Mouse Nervous System. Cell. 2018 Aug 9;174(4):999–1014.e22.

17. Zhou J, Yang C-X, Zhong J-Y, Wang H-B. Intrathecal TRESK gene recombinant adenovirus attenuates spared nerve injury-induced neuropathic pain in rats. Neuroreport. 2013 Feb;24(3):131–6.

18. Guo Z, Cao Y-Q. Over-expression of TRESK K(+) channels reduces the excitability of trigeminal ganglion nociceptors. PLoS ONE. 2014;9(1):e87029.

19. Zhou J, Yao S-L, Yang C-X, Zhong J-Y, Wang H-B, Zhang Y. TRESK gene recombinant adenovirus vector inhibits capsaicin-mediated substance P release from cultured rat dorsal root ganglion neurons. Mol Med Report. 2012 Apr;5(4):1049–52.

20. Lafrenière RG, Cader MZ, Poulin J-F, Andres-Enguix I, Simoneau M, Gupta N, Boisvert K, Lafrenière F, Mclaughlan S, Dubé M-P, Marcinkiewicz MM, Ramagopalan S, Ansorge O, Brais B, Sequeiros J, Pereira-Monteiro JM, Griffiths LR, Tucker SJ, Ebers G, Rouleau GA. A dominant-negative mutation in the TRESK potassium channel is linked to familial migraine with aura. Nat Med. 2010 Oct;16(10):1157–60.

21. Royal P, Andres-Bilbe A, Ávalos Prado P, Verkest C, Wdziekonski B, Schaub S, Baron A, Lesage F, Gasull X, Levitz J, Sandoz G. Migraine-Associated TRESK Mutations Increase Neuronal Excitability through Alternative Translation Initiation and Inhibition of TREK. Neuron. 2019 Jan 16;101(2):232–6.

22. Chae YJ, Zhang J, Au P, Sabbadini M, Xie G-X, Yost CS. Discrete change in volatile anesthetic sensitivity in mice with inactivated tandem pore potassium ion channel TRESK. Anesthesiology. 2010 Dec;113(6):1326–37.

23. Kollert S, Dombert B, Döring F, Wischmeyer E. Activation of TRESK channels by the inflammatory mediator lysophosphatidic acid balances nociceptive signalling. Sci Rep. 2015;5:12548.

24. Yang Y, Li S, Jin Z-R, Jing H-B, Zhao H-Y, Liu B-H, Liang Y-J, Liu L-Y, Cai J, Wan Y, Xing G-G. Decreased abundance of TRESK two-pore domain potassium channels in sensory neurons underlies the pain associated with bone metastasis. Sci Signal. American Association for the Advancement of Science; 2018 Oct 16;11(552):eaao5150.

25. Abbott FV, Franklin KB, Westbrook RF. The formalin test: scoring properties of the first and second phases of the pain response in rats. Pain. 1995 Jan;60(1):91–102.

26. Tjølsen A, Berge OG, Hunskaar S, Rosland JH, Hole K. The formalin test: an evaluation of the method. Pain. 1992 Oct;51(1):5–17.

27. McNamara CR, Mandel-Brehm J, Bautista DM, Siemens J, Deranian KL, Zhao M, Hayward NJ, Chong JA, Julius D, Moran MM, Fanger CM. TRPA1 mediates formalin-induced pain. Proc Natl Acad Sci USA. 2007 Aug 14;104(33):13525–30.

28. Fischer M, Carli G, Raboisson P, Reeh P. The interphase of the formalin test. Pain. 2013 Nov 27;155(3):511–21.

29. Alessandri-Haber N, Joseph E, Dina OA, Liedtke W, Levine JD. TRPV4 mediates pain-related behavior induced by mild hypertonic stimuli in the presence of inflammatory mediator. Pain. 2005 Nov;118(1-2):70–9.

30. Alessandri-Haber N, Yeh JJ, Boyd AE, Parada CA, Chen X, Reichling DB, Levine JD. Hypotonicity induces TRPV4-mediated nociception in rat. Neuron. 2003 Jul 31;39(3):497–511.

31. Yalcin I, Charlet A, Freund-Mercier M-J, Barrot M, Poisbeau P. Differentiating Thermal Allodynia and Hyperalgesia Using Dynamic Hot and Cold Plate in Rodents. The Journal of Pain. 2009 Jul;10(7):767–73.

32. Lolignier S, Gkika D, Andersson D, Leipold E, Vetter I, Viana F, Noël J, Busserolles J. New Insight in Cold Pain: Role of Ion Channels, Modulation, and Clinical Perspectives. J Neurosci. 2016 Nov 9;36(45):11435–9.

33. Bautista DM, Sigal YM, Milstein AD, Garrison JL, Zorn JA, Tsuruda PR, Nicoll RA, Julius D. Pungent agents from Szechuan peppers excite sensory neurons by inhibiting two-pore potassium channels. Nat Neurosci. 2008 Jul;11(7):772–9.

34. LaPaglia DM, Sapio MR, Burbelo PD, Thierry-Mieg J, Thierry-Mieg D, Raithel SJ, Ramsden CE, Iadarola MJ, Mannes AJ. RNA-Seq investigations of human post-mortem trigeminal ganglia. Cephalalgia. 2018 Apr;38(5):912–32.

35. Nguyen MQ, Wu Y, Bonilla LS, Buchholtz von LJ, Ryba NJP. Diversity amongst trigeminal neurons revealed by high throughput single cell sequencing. Obukhov AG, editor. PLoS ONE. Public Library of Science; 2017 Sep 28;12(9):e0185543–22.

36. Chiu IM, Barrett LB, Williams EK, Strochlic DE, Lee S, Weyer AD, Lou S, Bryman G, Roberson DP, Ghasemlou N, Piccoli C, Ahat E, Wang V, Cobos EJ, Stucky CL, Ma Q, Liberles SD, Woolf C. Transcriptional profiling at whole population and single cell levels reveals somatosensory neuron molecular diversity. Elife. 2014 Dec 19;3:e04660.

37. Li C-L, Li K-C, Wu D, Chen Y, Luo H, Zhao J-R, Wang S-S, Sun M-M, Lu Y-J, Zhong Y-Q, Hu X-Y, Hou R, Zhou B-B, Bao L, Xiao H-S, Zhang X. Somatosensory neuron types identified by high-coverage single-cell RNA-sequencing and functional heterogeneity. Cell Res. 2016 Jan;26(1):83–102.

38. Du X, Gao H, Jaffe D, Zhang H, Gamper N. M-type K +channels in peripheral nociceptive pathways. British Journal of Pharmacology. John Wiley & Sons, Ltd (10.1111); 2018;175(12):2158–72.

39. Castellanos A, Andres-Bilbe A, Bernal L, Callejo G, Comes N, Gual A, Giblin JP, Roza C, Gasull X. Pyrethroids inhibit K2P channels and activate sensory neurons. Pain. 2018 Jan;159(1):92–105.

40. Lu R, Lukowski R, Sausbier M, Zhang DD, Sisignano M, Schuh C-D, Kuner R, Ruth P, Geisslinger G, Schmidtko A. BKCa channels expressed in sensory neurons modulate inflammatory pain in mice. Pain. 2013 Dec 11;155(3):556–65.

41. Descoeur J, Pereira V, Pizzoccaro A, Francois A, Ling B, Maffre V, Couette B, Busserolles J, Courteix C, Noël J, Lazdunski M, Eschalier A, Authier N, Bourinet E. Oxaliplatin-induced cold hypersensitivity is due to remodelling of ion channel expression in nociceptors. EMBO Mol Med. 2011 May;3(5):266–78.

42. Zimmermann K, Leffler A, Babes A, Cendan CM, Carr RW, Kobayashi J-I, Nau C, Wood JN, Reeh PW. Sensory neuron sodium channel Nav1.8 is essential for pain at low temperatures. Nature. 2007 Jun 14;447(7146):855–8.

43. Story GM, Peier AM, Reeve AJ, Eid SR, Mosbacher J, Hricik TR, Earley TJ, Hergarden AC, Andersson DA, Hwang SW, McIntyre P, Jegla T, Bevan S, Patapoutian A. ANKTM1, a TRP-like channel expressed in nociceptive neurons, is activated by cold temperatures. Cell. 2003 Mar 21;112(6):819–29.

44. Yarmolinsky DA, Peng Y, Pogorzala LA, Rutlin M, Hoon MA, Zuker CS. Coding and Plasticity in the Mammalian Thermosensory System. Neuron. Elsevier; 2016 Dec 7;92(5):1079–92.

45. Memon T, Chase K, Leavitt LS, Olivera BM, Teichert RW. TRPA1 expression levels and excitability brake by KV channels influence cold sensitivity of TRPA1-expressing neurons. Neuroscience. 2017 Jun 14;353:76–86.

46. Madrid R, la Peña de E, Donovan-Rodriguez T, Belmonte C, Viana F. Variable threshold of trigeminal cold-thermosensitive neurons is determined by a balance between TRPM8 and Kv1 potassium channels. J Neurosci. 2009 Mar 11;29(10):3120–31.

47. Murthy SE, Loud MC, Daou I, Marshall KL, Schwaller F, Kühnemund J, Francisco AG, Keenan WT, Dubin AE, Lewin GR, Patapoutian A. The mechanosensitive ion channel Piezo2 mediates sensitivity to mechanical pain in mice. Sci Transl Med. American Association for the Advancement of Science; 2018 Oct 10;10(462):eaat9897.

48. Szczot M, Liljencrantz J, Ghitani N, Barik A, Lam R, Thompson JH, Bharucha-Goebel D, Saade D, Necaise A, Donkervoort S, Foley AR, Gordon T, Case L, Bushnell MC, Bönnemann CG, Chesler AT. PIEZO2 mediates injury-induced tactile pain in mice and humans. Sci Transl Med. American Association for the Advancement of Science; 2018 Oct 10;10(462):eaat9892.

49. Dhandapani R, Arokiaraj CM, Taberner FJ, Pacifico P, Raja S, Nocchi L, Portulano C, Franciosa F, Maffei M, Hussain AF, de Castro Reis F, Reymond L, Perlas E, Garcovich S, Barth S, Johnsson K, Lechner SG, Heppenstall PA. Control of mechanical pain hypersensitivity in mice through ligand-targeted photoablation of TrkB-positive sensory neurons. Nature Communications. Nature Publishing Group; 2018 Apr 24;9(1):1640.

50. Ranade SS, Woo S-H, Dubin AE, Moshourab RA, Wetzel C, Petrus M, Mathur J, Bégay V, Coste B, Mainquist J, Wilson AJ, Francisco AG, Reddy K, Qiu Z, Wood JN, Lewin GR, Patapoutian A. Piezo2 is the major transducer of mechanical forces for touch sensation in mice. Nature. 2014 Dec 4;516(7529):121–5.

51. Beaulieu-Laroche L, Christin M, Donoghue A, Agosti F, Yousefpour N, Petitjean H, Davidova A, Stanton C, Khan U, Dietz C, Faure E, Fatima T, MacPherson A, Ribeiro-da-Silva A, Bourinet E, Blunck R, Sharif-Naeini R. TACAN is an essential component of the mechanosensitive ion channel responsible for pain sensing. bioRxiv. 2018 Jan 1;:338673.

52. Cavanaugh DJ, Lee H, Lo L, Shields SD, Zylka MJ, Basbaum AI, Anderson DJ. Distinct subsets of unmyelinated primary sensory fibers mediate behavioral responses to noxious thermal and mechanical stimuli. Proc Natl Acad Sci U S A. 2009 Jun 2;106(22):9075–80.

53. Vriens J, Nilius B, Voets T. Peripheral thermosensation in mammals. Nat Rev Neurosci. 2014 Jul 23;15(9):573–89.

54. Julius D. TRP channels and pain. Annu Rev Cell Dev Biol. 2013;29:355–84.

55. Callejo G, Giblin JP, Gasull X. Modulation of TRESK background K+ channel by membrane stretch. Ceña V, editor. PLoS ONE. 2013;8(5):e64471.

56. Bhattacharya MRC, Bautista DM, Wu K, Haeberle H, Lumpkin EA, Julius D. Radial stretch reveals distinct populations of mechanosensitive mammalian somatosensory neurons. Proc Natl Acad Sci U S A. 2008 Dec 16;105(50):20015–20.

57. Chaplan SR, Bach FW, Pogrel JW, Chung JM, Yaksh TL. Quantitative assessment of tactile allodynia in the rat paw. J Neurosci Methods. 1994 Jul;53(1):55–63.

58. Brenner DS, Golden JP, Gereau RW IV. A novel behavioral assay for measuring cold sensation in mice. PLoS ONE. Public Library of Science; 2012;7(6):e39765.

59. Yalcin I, Megat S, Barthas F, Waltisperger E, Kremer M, Salvat E, Barrot M. The sciatic nerve cuffing model of neuropathic pain in mice. J Vis Exp. 2014 Jul 16;(89).

60. Callejo G, Castellanos A, Castany M, Gual A, Luna C, Acosta MC, Gallar J, Giblin JP, Gasull X. Acid-sensing ion channels detect moderate acidifications to induce ocular pain. Pain. 2015 Mar;156(3):483–95.

61. Le Pichon CE, Chesler AT. The functional and anatomical dissection of somatosensory subpopulations using mouse genetics. Front Neuroanat. Frontiers; 2014;8:21.

