## Supplementary material for "TRESK background K^+^ channel deletion selectively uncovers enhanced mechanical and cold sensitivity": Suppl fig and table

Dynamic hot plate

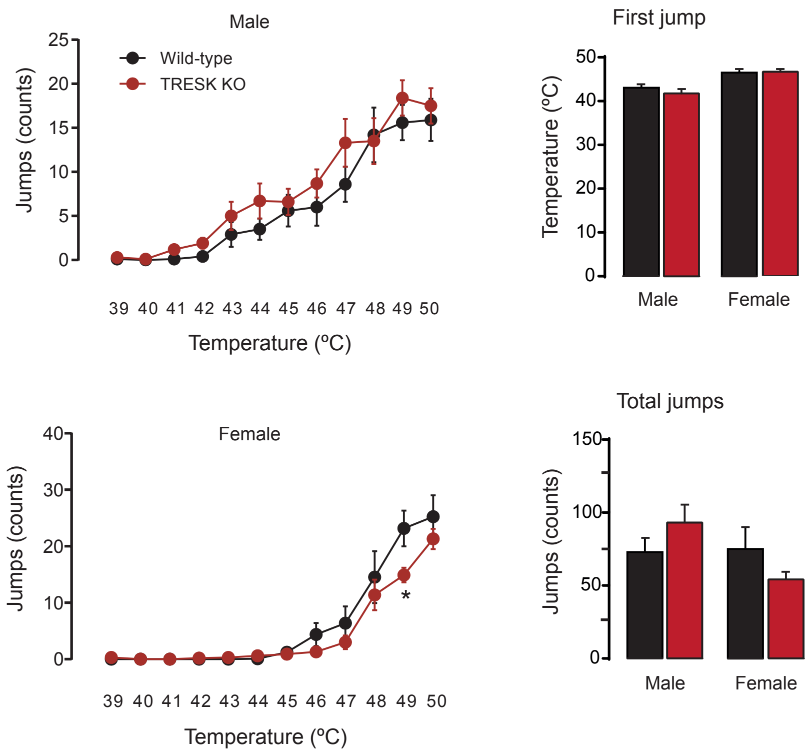


**Supplementary Figure 1: Dynamic hot plate**
In contrast to the conventional hot plate, the dynamic hot plate allows the testing of a wide range of temperatures. Plate temperature was ramped from 30ºC to 50ºC at a 1ºC/min rate. To determine the temperature that is perceived as noxious for mice and quantify pain-related behaviors, the number of jumps at each temperature was scored. *Left:* number of jumps elicited at each temperature in male (*top*) and female (*bottom*) in wild-type and TRESK KO mice (n=11-15 animals per group). Only a significant difference was obtained at 49ºC ($p<0.05$, unpaired t-test) between wild-type and TRESK KO females. *Right, top:* mean temperature at which animals did the first jump (threshold). *Right. bottom:* Total number of jumps for male and female mice in the whole range of temperatures. Values for temperatures between 30 and 38ºC are not shown in the plots since were probably detected as non-noxious and did not produce any observable response.

**Supplementary Table 1**

Primer sequences used for quantitative real-time PCR

| Target | Common name | PCR primer sequence (5'-3') | |
|---|---|---|---|
| Kcnk18 | TRESK | For | CTCTCTTCTCCGCTGTCGAG |
| | | Rev | AAGAGAGCGCTCAGGAAGG |
| Kcnk2 | TREK-1 | For | CACTGTGAGTTTTGCACATGG |
| | | Rev | GGGACTGGACTTTTTCTGAATC |
| Kcnk10 | TREK-2 | For | GCAGCTTTCCCTTAGACCAG |
| | | Rev | CCAGGGACATTCATTTTGGA |
| Kcnk4 | TRAAK | For | CATCCAAAAAGCCTTCCAGA |
| | | Rev | ATTTGGCAACCACTGGACTC |
| Trpa1 | TRPA1 | For | GCAGGTGGAACTTCATACCAACT |
| | | Rev | CACTTTGCGTAAGTACCAGAGTGG |
| Trpv1 | TRPV1 | For | CCCATTGTGCAGATTGAGCAT |
| | | Rev | TTCCTGCAGAAGAGCAAGAAGC |
| Gapdh | GAPDH | For | ATGTGTCCGTCGTGGATCTGA |
| | | Rev | GCTGTTGAAGTCGCAGGAGAC |
